# Subjective estimates of uncertainty during gambling and impulsivity after subthalamic deep brain stimulation for Parkinson’s disease

**DOI:** 10.1101/477364

**Authors:** Saee Paliwal, Philip E. Mosley, Michael Breakspear, Terry Coyne, Peter Silburn, Eduardo Aponte, Christoph Mathys, Klaas E. Stephan

**Author notes:** Correspondence to: Dr Philip E Mosley, Neurosciences Queensland, Level 1, St Andrew’s Place, 33 North Street, Spring Hill, Queensland, 4000, Australia. These authors contributed equally to the work (Saee Paliwal, Philip E. Mosley).

## Abstract

Subthalamic deep brain stimulation (STN-DBS) for Parkinson’s disease may modulate chronometric and instrumental aspects of choice behaviour, including motor inhibition, decisional slowing, and value sensitivity. However, it is not well known whether STN-DBS affects more complex aspects of decision-making, such as the influence of subjective estimates of uncertainty on choices. In this study, thirty-eight participants with Parkinson’s disease played a virtual casino prior to subthalamic DBS (whilst ‘*on*’ medication) and again, three-months postoperatively (whilst ‘*on*’ stimulation). At the group level, there was a small but statistically significant postoperative decrease in impulsivity, as quantified by the Barratt Impulsiveness Scale (BIS). The gambling behaviour of participants (bet increases, slot machine switches and double or nothing gambles) was associated with this self-reported measure of impulsivity. However, there was a large variance in outcome amongst participants, and we were interested in whether individual differences in subjective estimates of uncertainty (specifically, volatility) were related to differences in pre- and postoperative impulsivity. To examine these individual differences, we fit a computational model (the Hierarchical Gaussian Filter, HGF), to choices made during slot machine game play as well as a simpler reinforcement learning model based on the Rescorla-Wagner formalism. The HGF was superior in accounting for the behaviour of our participants, suggesting that participants incorporated beliefs about environmental uncertainty when updating their beliefs about gambling outcome and translating these beliefs into action. A specific aspect of subjective uncertainty, the participant’s estimate of the tendency of the slot machine’s winning probability to change (volatility), increased subsequent to DBS. Additionally, the decision temperature of the response model decreased post-operatively, implying greater stochasticity in the belief-to-choice mapping of participants. Model parameter estimates were significantly associated with impulsivity; specifically, increased uncertainty was related to increased postoperative impulsivity. Moreover, changes in these parameter estimates were significantly associated with the maximum post-operative change in impulsivity over a six month follow up period. Our findings suggest that impulsivity in persons with Parkinson’s disease may be influenced by subjective estimates of uncertainty (environmental volatility) and implicate a role for the subthalamic nucleus in the modulation of outcome certainty. Furthermore, our work outlines a possible approach to characterising those persons who become more impulsive after subthalamic DBS, an intervention in which non-motor outcomes can be highly variable.

## 2 Introduction

The subthalamic nucleus (STN) is a subcortical nucleus of central pathophysiological relevance for Parkinson’s disease. In Parkinson’s disease, STN neurons display abnormal patterns of burst firing (Vila *et al.*, 2000) and low-frequency synchronisation of local field potentials (Brown *et al.*, 2001). By modulating this pathological network activity, subthalamic deep brain stimulation (STN-DBS) improves motor symptoms such as bradykinesia, tremor and rigidity (Schuepbach *et al.*, 2013). However, STN-DBS has also been linked to neuropsychiatric symptoms, particularly impulsivity (Jahanshahi *et al.*, 2015; Mosley and Marsh, 2015), an issue of substantial clinical importance. The STN is a second input station to the basal ganglia, receiving direct cortical projections from the frontal lobe (the ‘*hyperdirect*’ pathway) (Nambu *et al.*, 2002). This route permits basal ganglia inhibitory tone to be directly modulated by cortical regions. Functional and structural brain imaging support the role of this pathway in motor inhibition (Aron *et al.*, 2007; Rae *et al.*, 2015). Following STN-DBS, participants with Parkinson’s disease make commission errors (in which participants execute an erroneous action) (Hershey *et al.*, 2004) and take longer to cancel an action (Obeso *et al.*, 2013), suggesting an impairment in action restraint. Errors in the Stroop (Witt *et al.*, 2004) and random number generation tasks (Thobois *et al.*, 2007) suggest increased sensitivity to cognitive interference and impaired task-switching. When faced with decisional conflict, persons with subthalamic DBS speed up rather than slow down their decision-making (Cavanagh *et al.*, 2011; Frank *et al.*, 2007). These findings suggest a role for the STN in ‘*braking*’ cognitive-associative circuits in the basal ganglia. It is not clear, however, whether such chronometric aspects of decision-making are sufficient to explain the complex picture of impulsivity after STN-DBS (Florin *et al.*, 2013). For example, subthalamic DBS may modulate contingencies in reinforcement learning (Seymour *et al.*, 2016; Wagenbreth *et al.*, 2015), suggesting a role for the STN in valuation.

A potential computational mechanism underlying impulsivity is the estimation of uncertainty. If the longer-term outcomes of actions are not (or do not seem to be) predictable, prospective thinking may be replaced by seeking immediately available outcomes, a policy that would manifest behaviourally as impulsivity (Patton *et al.*, 1995). More specifically, subjective uncertainty about environmental dynamics, or the longer-term consequences of actions, has been associated with a tendency to reduce reflection and long-term planning, and favour short-term over long-term outcomes (Averbeck *et al.*, 2014; FitzGerald *et al.*, 2015; Paliwal *et al.*, 2014). For example, consider deciding how to invest a sum of money in order to maximise profit. In a dependable economic setting with a reliable government, low unemployment, steadily rising house prices and a stable stock market, one might be confident that a long-term investment in property, business or shares would pay off highly. However, if the long-term outlook was unpredictable, with the possibility of a military coup, high unemployment, crashing property market and volatile stocks, one might decide to spend the money now and enjoy its worth in the short-term. This connection between uncertainty and impulsivity is also seen in Parkinson’s disease. Persons with Parkinson’s disease and impulse control behaviours (ICBs) ‘*jump to conclusions*’ in an information collection task (the beads task) more quickly than participants without ICBs, a finding that relates informational uncertainty to impulsivity (Djamshidian *et al.*, 2012). A computational modelling study of behaviour across three tasks commonly used to probe impulsivity (information sampling, temporal discounting and novelty bias) suggested that Parkinson’s disease participants with ICBs are more uncertain about the relationship between possible actions and future rewards than patients without ICBs (Averbeck *et al.*, 2013). Based on these findings, we hypothesised that changes in impulsivity after STN-DBS may also relate to estimates of environmental uncertainty about future rewards.

A paradigmatic approach to inference and learning under uncertainty uses Bayes’ theorem to understand how prior knowledge (represented as a probability distribution known as the *prior*) is combined with new information from the environment (the *likelihood*) in order to update beliefs (the *posterior*). To obtain the posterior, a Bayesian agent inverts a ‘*generative*’ model (that describes how noisy sensory data result from environmental states); this corresponds to perception. Inferring environmental states from noisy sensory data allows the agent to plan actions that take into account the uncertainty of the environment (Daunizeau *et al.*, 2010). Human behaviour often closely resembles those of Bayesian agents, for example, during low level sensory processing (Petzschner *et al.*, 2015; Weiss *et al.*, 2002), sensorimotor learning (Kording and Wolpert, 2004; Wolpert *et al.*, 1995) and higher-level reasoning (Tenenbaum *et al.*, 2006), although approximations to ideal Bayesian inference are likely required for most domains of cognition (Friston, 2009; Griffiths *et al.*, 2010; Tenenbaum *et al.*, 2011).

Critically, a Bayesian perspective can accommodate multiple forms of uncertainty, beyond sensory noise. For example, the agent’s environment might change over time. In order to account for this environmental uncertainty (or volatility), Bayesian agents are able to modulate the rate at which they learn (update their beliefs). This learning rate can be linked to an agent’s encoding of volatility (Behrens *et al.*, 2007; Mathys *et al.*, 2011; Mathys *et al.*, 2014). For instance, in more volatile environments, estimates of uncertainty (and thus learning rate) should be higher so that more emphasis is given to very recent information; at the same time, predicting the longer-term consequences of actions becomes more difficult. Additionally, uncertainty around how to best translate an uncertain belief about the environment into the selection of an action that will eventually lead to a reward is yet another source of noise that could contribute to observed stochasticity in behaviour. This link between uncertainty and decision-making may be of crucial importance for impulsivity (Averbeck *et al.*, 2013). Furthermore, individual differences in approximate Bayesian inference plausibly contribute to inter-individual variability in behaviour. Such differences can be quantified using models with participant-specific parameters concerning, for instance, the estimation of environmental volatility (Vossel *et al.*, 2014), or the formation of unusually confident or ‘*precise*’ beliefs (Schwartenbeck *et al.*, 2015).

In prior work, we found that participant-specific parameter estimates encoding different aspects of uncertainty (such as an estimate of environmental uncertainty, the tonic volatility *ω*) related to a clinical measure of impulsivity in a cohort of healthy individuals (Paliwal *et al.*, 2014). This finding supported a mechanistic understanding of impulsivity from the perspective of uncertainty. Based on these results, we sought to assess whether similar relationships between computational measures of uncertainty and impulsivity were observed in a population of persons with Parkinson’s disease undertaking subthalamic DBS. Moreover, given the central role of this brain region in decision-making, we investigated whether receiving subthalamic DBS was associated with changes in the encoding of uncertainty that were connected to increases in impulsivity in certain persons. Finally, we assessed whether a perioperative computational characterisation of subjective aspects of uncertainty (estimates of environmental volatility and stochasticity of belief-response mappings) would be associated with longitudinal changes in impulsivity during clinical follow up. This latter question is clinically important as the non-motor outcomes from subthalamic DBS can be varied. Some centres advocate for the use of DBS to address impulsivity and compulsivity amongst persons with Parkinson’s disease on dopamine agonist medication, as subthalamic DBS allows these agents to be substantially reduced or even withdrawn (Lhommee *et al.*, 2012). However, other centres report the emergence of harmful impulsivity subsequent to DBS, in persons with no prior history of clinically-significant psychiatric symptoms (Amami *et al.*, 2015; Halbig *et al.*, 2009; Lim *et al.*, 2009; Mosley *et al.*, 2018b; Smeding *et al.*, 2007). At present, there is little evidence to guide the identification of surgical candidates at risk of postoperative impulsivity (Mosley and Marsh, 2015; Voon *et al.*, 2006).

In this analysis, we employed a similar computational framework to that previously reported (Paliwal *et al.*, 2014), applying a hierarchical Bayesian model (the Hierarchical Gaussian Filter, HGF) to behavioural data from thirty-eight participants with Parkinson’s disease who played a virtual casino before and after subthalamic DBS. By allowing participants to vary their bet size, switch between slot machines and place double-or-nothing bets, we could estimate how participants not only inferred the trial-by-trial probability of winning, but also updated higher-order beliefs about the fluctuations (volatility) of a slot machine’s winning probability. Similar to the rationale outlined in prior work,(Paliwal *et al.*, 2014) we believe that a naturalistic paradigm engenders increased behavioural engagement, allowing us to quantify behaviour that has a higher fidelity to ‘*real world*’ impulsivity. Additionally, model-based estimates derived from the computational framework may afford us an individual profile of how each participant represented (and responded to) environmental uncertainty. We assessed these findings against standard measures of impulsivity derived from clinical assessment and questionnaires, focusing our attention on self-reported impulsivity as measured by the Barratt Impulsiveness Scale (BIS). Our computational analysis examined how DBS changes the manner in which persons with Parkinson’s disease engage in Bayesian belief updating, and whether changes in participant-specific estimates of uncertainty before and after DBS relate to changes in impulsivity.

## 3 Methods

### 3.1 Participants

Thirty-eight participants with Parkinson’s disease undertaking STN-DBS were consecutively recruited at the Asia-Pacific Centre for Neuromodulation in Brisbane, Australia. All participants met the UK Brain Bank criteria for Parkinson’s disease (Hughes *et al.*, 1992). No participants met the Movement Disorder Society criteria for dementia (Emre *et al.*, 2007). The Parkinson’s disease subtype and the Hoehn and Yahr stage at device implantation was recorded (Hoehn and Yahr, 1967). Participants underwent bilateral implantation of Medtronic 3389 or Boston Vercise electrodes in a single-stage procedure. Stimulation was commenced immediately using microelectrode recording data to identify the optimal contact. Further contact testing took place over the following week as an inpatient, with participants returning to the DBS centre following discharge for further stimulation titration, guided by residual motor symptoms. Further details have previously been reported (Mosley *et al.*, 2018a; Mosley *et al.*, 2018c).

### 3.2 Neuropsychiatric Assessment

Impulsivity amongst participants was assessed with patient and clinician-rated instruments prior to STN-DBS and subsequently two-weeks, six-weeks, thirteen-weeks and twenty-six-weeks postoperatively (Figure 1). A range of measures were obtained, to account for the fact that impulsivity is not a unitary construct. These included the Barratt Impulsiveness Scale 11 (BIS) and second-order factors attentional, motor and non-planning (Patton *et al.*, 1995); the Questionnaire for Impulsive-Compulsive disorders in PD Rating Scale (QUIP-RS) (Weintraub *et al.*, 2012); the delay discounting task (Kirby *et al.*, 1999); the Excluded letter fluency task (ELF) (Shores *et al.*, 2006); and the Hayling test (Burgess *et al.*, 1997).

**Figure 1.**
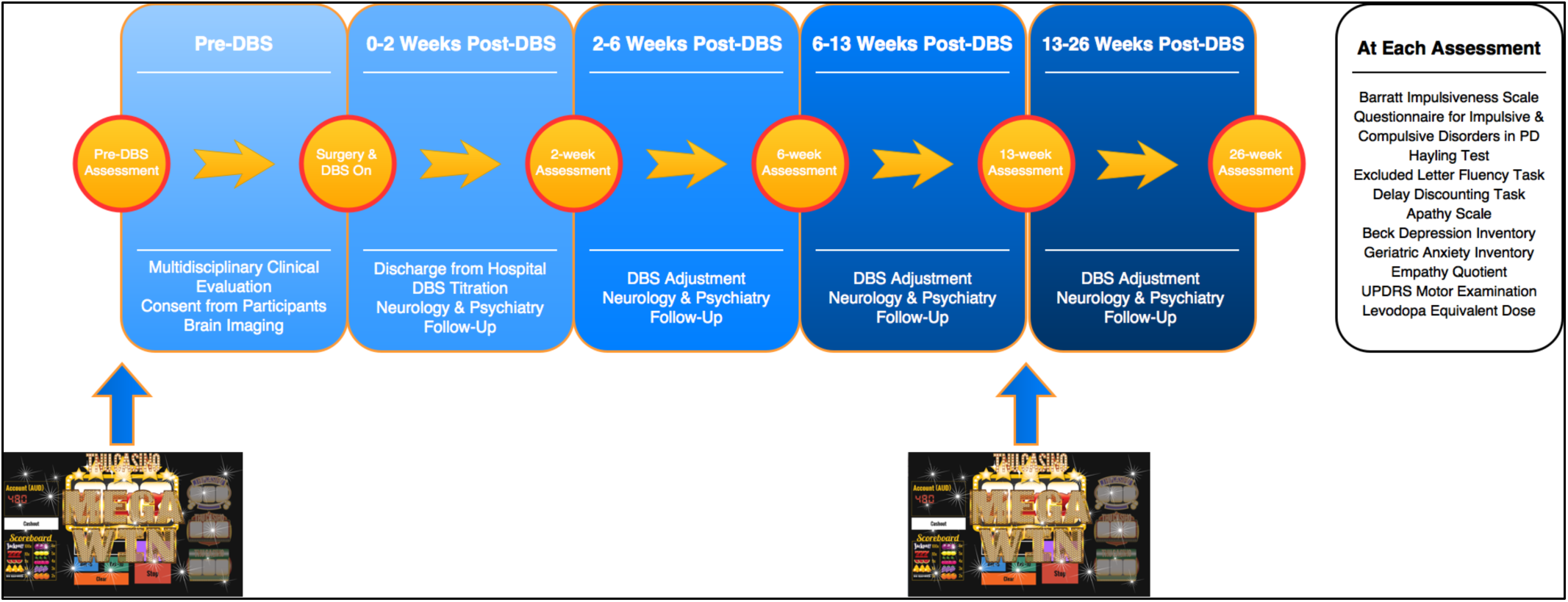
Overview of study timeline. Persons with Parkinson’s disease were screened in a multidisciplinary clinic by a movement disorders neurologist, neurosurgeon, psychiatrist, and rehabilitation specialist prior to selection for subthalamic DBS. Prior to surgery, surgical candidates who consented to participate were assessed with a battery of neuropsychiatric instruments (see box). These instruments were repeated at 2-weeks, 6-weeks, 13-weeks and 26-weeks postoperatively. This methodology of iterative assessments was designed to capture fluctuations in neuropsychiatric symptoms as electrical stimulation was titrated and dopaminergic medication was reduced. The slot machine gambling paradigm was undertaken by participants prior to neurosurgery and at 13-weeks postoperatively.

#### 3.2.1 Barratt Impulsiveness Scale

The BIS is one of the most widely used instruments for assessing trait impulsivity and is often the gold standard instrument against which other measures are compared. It is a thirty-item self-report questionnaire assessing the prevalence of impulsive behaviours. Respondents must rate each item (e.g. ‘*I act on the spur of the moment*’) from 1 to 4 according to the frequency of occurrence (i.e. rarely / never; occasionally; often; always / almost always). Higher scores indicate greater impulsivity. The mean BIS score is significantly greater in Parkinson’s disease participants with ICBs compared to non-impulsive persons with Parkinson’s disease (Isaias *et al.*, 2008; Voon *et al.*, 2007), suggesting that this instrument also has construct validity in the assessment of impulsivity in Parkinson’s disease.

#### 3.2.2 Questionnaire for Impulsive-Compulsive disorders in PD

The QUIP-RS is a twenty-eight-item self-report questionnaire assessing the prevalence of ICBs including compulsive spending, hypersexuality, pathological gambling, binge eating, hobbyism, punding and dopamine dysregulation. Respondents much rate each item (e.g. ‘*Do you have urges or desires for the following behaviours that you feel are excessive or cause you distress?*’) from 0 (never) to 4 (very often). The total sum obtained for each compulsive behaviour indicates the current severity of that behaviour. The instrument was designed for use in the Parkinson’s disease population and was previously validated against a semi-structured clinical interview (Weintraub *et al.*, 2012).

#### 3.2.3 Hayling Test

The Hayling Test is a sentence completion task, during which participants must insert a nonsense word at the end of a sentence, inhibiting the pre-potent stimulus to complete the sentence with a word that makes sense. The test assesses the construct of inhibition and is sensitive to frontal lobe dysfunction. For example, in the sentence: ‘*the whole town came to hear the Mayor*…’ a correct response could be ‘*banana*’. Participants would be penalised for completing the sentence with the clearly related words ‘*speak*’, or ‘*talk*’ (referred to as category A errors), as well as with words that are only partially connected such as ‘*explode*’ (referred to as category B errors). Parkinson’s disease participants make more category A and B errors than controls on this task (O’Callaghan *et al.*, 2013a; Obeso *et al.*, 2011).

#### 3.2.4 Excluded Letter Fluency Task

The ELF is an additional measure of inhibitory control. Participants are given three trials of ninety seconds to produce as many words as possible that do not contain a specified vowel. Words must be longer than three letters and cannot be proper nouns or derivations of the same word stem. Scoring includes an overall correct total, the number of rule violations and the number of word repetitions. In a sample of fifty persons with Parkinson’s disease, the number of rule violations was previously shown to be significantly greater compared to age-matched controls and was highly correlated with anatomical changes in brain regions implicated in inhibition (O’Callaghan *et al.*, 2013b).

#### 3.2.5 Delay Discounting Task

An assessment of delay aversion, the tendency to prefer sooner, smaller rewards over those that are larger but temporally more distant. The task was designed to assess impulsivity in individuals with substance use disorders; behaviours that share face validity with the impulse-control disorders (ICBs) observed in a subset of persons with Parkinson’s disease. Participants are presented with a series of twenty-seven choices between an amount of money distributed immediately and a larger sum after a specified delay. After the task is complete, participants have the opportunity to win the amount of money they have chosen in a choice selected at random, either immediately or after a delay, depending on the choice they have made. Subsequently, the pattern of choices is analysed to calculate a discount parameter, or the point of indifference between delayed and immediate rewards for a given sum. Individuals with greater delay aversion have a higher discount parameter. The extent of delay aversion was previously shown to be greater amongst persons with Parkinson’s disease than healthy controls (Milenkova *et al.*, 2011), as well as being greater amongst Parkinson’s disease participants with ICBs than non-impulsive persons with Parkinson’s disease (Housden *et al.*, 2010; Voon *et al.*, 2011).

Additional neuropsychiatric symptoms were captured with the Beck Depression Inventory II (BDI) (Beck *et al.*, 1961); the Empathy Quotient (EQ) (Baron-Cohen and Wheelwright, 2004); the Geriatric anxiety inventory (GAI) (Pachana *et al.*, 2007) and the Apathy Scale (Starkstein *et al.*, 1992). For each self-report scale, participants were instructed to refer to ‘*the last two weeks*’, in order to obtain a measurement of current ‘*state*’. At each visit, Parkinson’s disease motor symptoms were assessed using the UPDRS Part III motor examination (Goetz *et al.*, 2007). Dopaminergic medication was recorded and converted to a levodopa-equivalent daily dose (LEDD) value (Evans *et al.*, 2004).

### 3.3 Design and Setting

Participants completed the experimental task prior to DBS and at thirteen-weeks post-DBS. Participants were ‘*on*’ medication and stimulation for all assessments. We opted against a counterbalanced ‘*off*’ and ‘*on*’ DBS assessment at the same visit for several reasons. First, our aim was to provide a naturalistic insight into the subtle behavioural changes that emerge as patients transition from dopaminergic therapies to subthalamic stimulation; changes in levodopa equivalent daily dose were included as co-variates in our analyses. Second, our experience is that many patients would not tolerate the DBS ‘*off*’ state without severe discomfort. Thirdly, despite allowing DBS washout, plastic network effects of chronic DBS may persist and contaminate findings in an on-off design.

### 3.4 Virtual Casino

We employed a modified version of an established slot machine gambling paradigm validated in healthy controls (Paliwal *et al.*, 2014). The task was designed to have standard features normally attributed to slot machines (colours, sounds, banners), and its features were designed to mirror the specifications of Swiss and German slot machines (Figure 2). Due to its game-like feel, the task successfully elicits impulsive, risk-taking and exploratory behaviour in participants. Task behaviour has been previously shown to correlate with standard measures of impulsivity (i.e. the BIS). Players began the slot machine with 2000 AUD in their account, and played through 100 trials. The trajectory of win-loss outcomes was predetermined, ensuring that participants’ experience of rewards and losses were comparable in order and quantity. The trajectory results in a positive outcome (net winnings) for most participants. At the end of the task, participants were awarded up to 30 AUD in real money based on the size of these virtual winnings.

**Figure 2.**
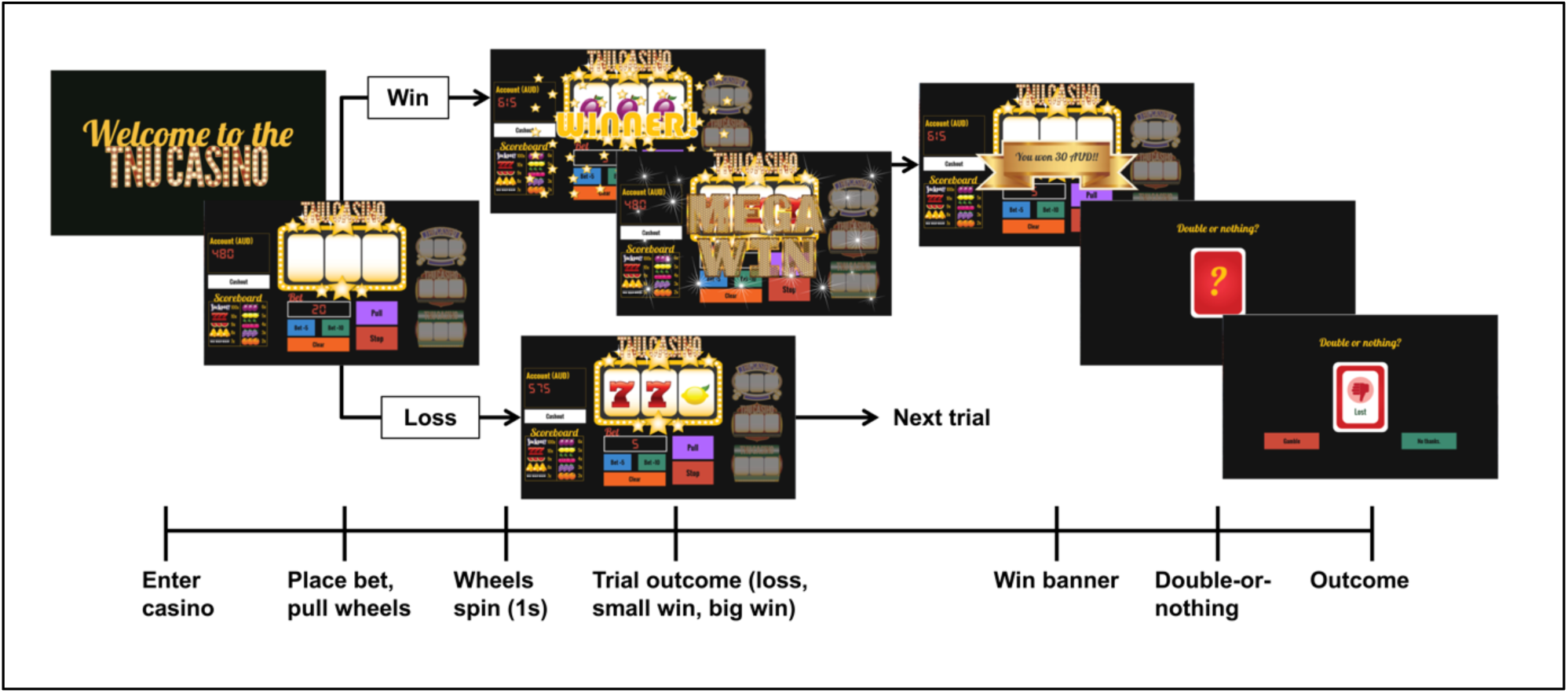
Slot machine gambling paradigm. The task consists of 100 trials. On every trial, players are able to place a bet of unlimited magnitude, switch slot machines or *‘cash out*’, exiting the casino and returning again on another virtual ‘*day*’. The overall win probability is 25 %, with wins split into big wins and small wins. The two possible types of losses are near-misses, in which the first two wheels are the same and the third is different (i.e. AAB) or a true loss, in which all the wheels are different (i.e. ABC). Game play proceeds as follows. Each trial begins with the slot machine main screen loading, displaying the player’s account value. The player then places a continuous-valued bet amount, incremented in units of 5 or 10 AUD. After the player has placed a bet, he or she presses the ‘*Pull*’ button and watches as the wheels begin to spin. At any point, the player has the ability to press the ‘*Stop*’ button, ending the trial and subsequently revealing the outcome of the three wheels. Unbeknownst to the participant, pressing the stop button has no effect on the trial outcome. If the stop button is not pressed, the trial times out after 5 seconds, and the player sees the outcome of the first, second and third wheel sequentially. On trials in which the outcome is a win, there are ten possible reward grades (or multiples of the bet amount). After every win trial, players are offered a possible ‘*double-up*’ option, during which players are given 3 seconds to decide whether or not to engage in a ‘*double-or-nothing*’ option, thereby risking his or her entire win amount. If the player elects to engage in this gamble, a card flips over revealing the result, and subjects are taken to the next trial. If the player does nothing, or decides not to gamble, he or she is taken to the next trial. For each loss trial, players are taken directly to the beginning of the next trial. Again, the trajectory of win-loss outcomes is fixed, ensuring comparable inference upon perceptual and response parameters across participants.

The task began with five training trials, after which the participant played through the main task, consisting of 100 trials. Only data from the main task were used for further analysis. For the main task, the win probability was 25 %, with wins split into big wins (12 % of trials) and small wins (88 % of trials). Players won when all three wheels showed the same symbols (e.g. all three wheels display an apple image). There were two possible types of losses. The first was a near-miss, in which the first two wheels of the slot machine displayed the same symbol, and the third was different (e.g. cherry, cherry, apple). The second was a true loss, in which all the wheels displayed different images (e.g. cherry, apple, orange).

Game play proceeded as follows: at the onset of each trial, the main screen loaded, displaying the player’s account value. Players were then able to execute one of the following actions: place a bet (of unlimited magnitude – by loading the machine in increments of 5-10 AUD), switch slot machines, or ‘*cash out’*, which involved ‘*exiting*’ the casino and returning again on another virtual ‘*day*’. If the player chose to bet, after loading the machine, they pressed the ‘*Pull*’ button and watched as the wheels begin to spin. The player had the option of pressing the ‘*Stop*’ button at any time during wheel spin, ending the trial and subsequently revealing the outcome of the three wheels, Pressing the stop button had no effect on the trial outcome; though this was not told to the participant. In the absence of the stop button being pressed, the trial timed out after five seconds, and revealed the outcome to the player, with the first, second and third wheel stopping sequentially. For winning trials, there were ten possible reward amounts. Each possible reward was called a reward grade, and indicated a different multiple of the bet size placed (e.g. reward grade 1 indicated a reward amount that was double the bet amount placed). After each win trial, the player is offered a ‘*double-up*’ option, during which they were given three seconds to decide whether or not to engage in a ‘*double-or-nothing*’ option. The double-or-nothing option had two possible outcomes: if the player won the double-or-nothing gamble, they doubled their win amount from that trial, if they lost the double-or-nothing option, they lost their entire win amount from the corresponding trial. If the player did nothing, or decided not to gamble, they are taken to the next trial. For losses, players were taken directly to the beginning of the next trial.

In the context of the analyses presented, this version of the slot machine deviated from the version presented in Paliwal *et al.* (2014) in several important ways: here, players were given the ability to place unlimited bet sizes with the ability to increase their bets in increments of 5 or 10 AUD; there were no ‘*fake win*’ results, simply wins and losses; the task was considerably (fifty percent) shorter; and finally, the task was aesthetically remodelled in order to be more naturalistic.

Each trial in the game followed a pre-programmed result sequence consisting of the following trial types:

i. big wins: top three out of 10 reward grades
ii. small wins: lower 7 out of 10 reward grades
iii. near misses: when the same symbol appeared in wheel 1 and two, and a different symbol appeared in wheel 3
iv. true losses (all three wheels showed different symbols).

After all win trials, a participant was allowed to engage in a secondary double-or-nothing gamble, and had three seconds to decide whether or not to do so. A more detailed trial breakdown across the various types of wins and losses is listed below:

i. 25 out of 100 trials were wins
ii. 3 out of 100 trials were big wins
iii. 22 out of 100 trials were small wins
iv. 22 out of 100 trials were near misses
v. 52 out of 100 trials were full losses.
vi. All win trials contained a double-up option

### 3.5 Computational Modelling

#### 3.5.1 The Hierarchical Gaussian Filter (HGF)

The HGF is a hierarchical Bayesian model (Mathys *et al.*, 2011; Mathys *et al.*, 2014) (Figure 3) where each level of the hierarchy encodes distributions of environmental variables (in ascending complexity) that evolve as Gaussian random walks. The HGF is an extension of the model presented in Behrens *et al.* (2007) and describes an agent whose learning rate is a function of his or her uncertainty. In the HGF, an agent is assumed not only to represent current environmental contingencies, but also to track how these contingencies change over time (volatility), and to what degree volatility itself is constant (tonic volatility) or may change in time (phasic volatility). Importantly, the agent modelled in the HGF employs these representations to make predictions about emerging environmental fluctuations and future sensory feedback. Furthermore, the agent is able to encode the precision of each prediction and use these precision estimates to scale trial-wise updates of beliefs about the environment and its statistical structure. Each level of the HGF is coupled such that higher states determine how quickly the next lower state evolves, with the lowest hierarchical level representing sensory events.

**Figure 3.**
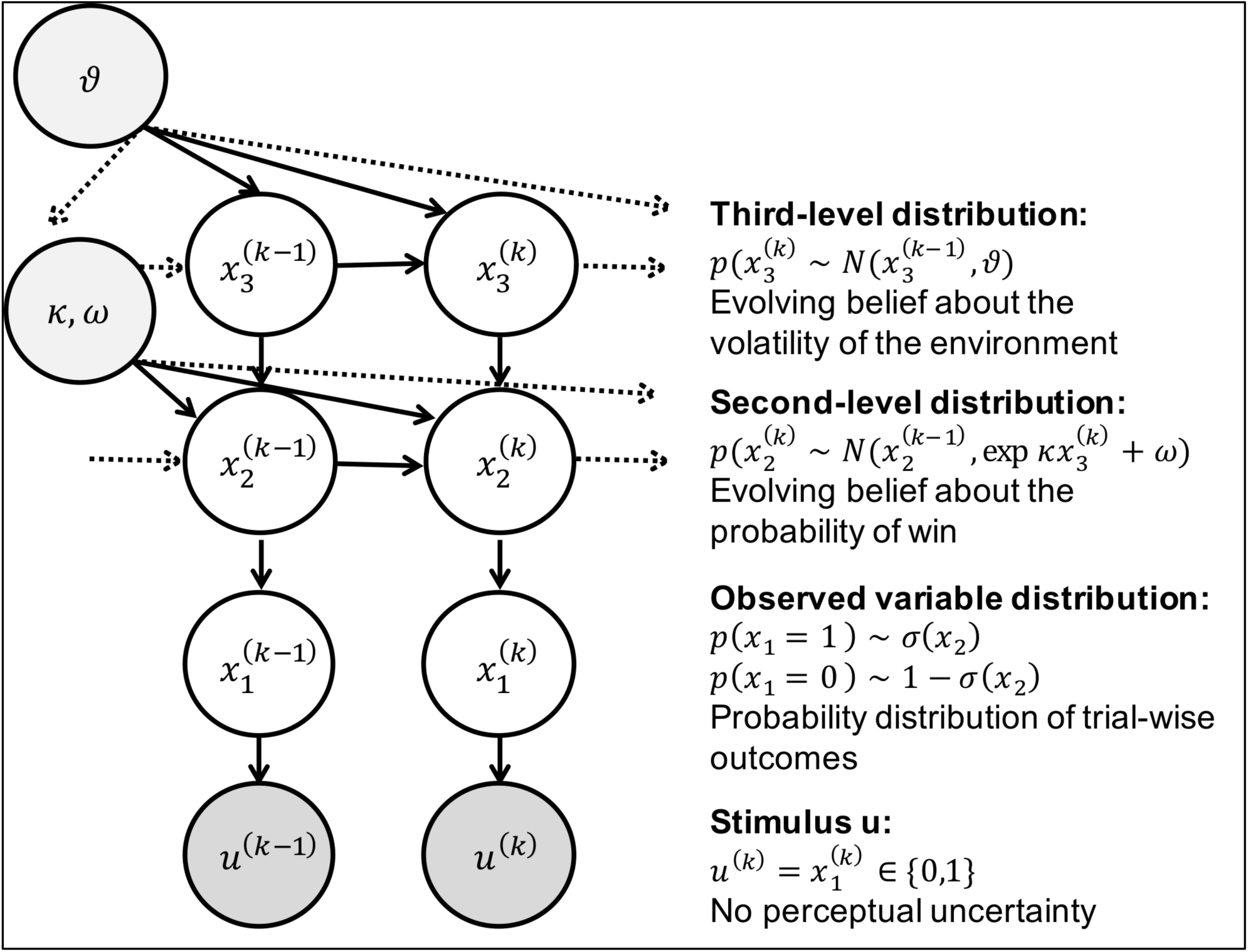
The Hierarchical Gaussian Filter (HGF) *u*^*(k)*^ represents binary observations (true wins=1, and losses=0, in the case of the slot machine). Binary inputs are represented on the first level, 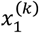 via a Bernoulli distribution, around the probability of win or loss, 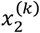. In turn, 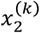 is modelled as a Gaussian random walk, whose step-size is governed by a combination of 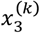, via coupling parameter *k*, and a tonic volatility parameter 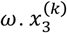 also evolves as a Gaussian random walk over trials, with step size *ϑ* (meta-volatility). In this investigation, after observing trial-wise outcomes (win or lose), the gambler updates her belief about the probability of win on a given trial 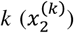, as well as how swiftly that slot machine is moving between being ‘*hot*’ (high probability of win) or ‘*cold*’ (low probability of win) 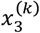. On any trial, the ensuing beliefs then provide a basis for the gambler’s response, which may be to increase the bet size, ‘*double up*’ after a win, switch to a new slot machine or leave the casino.

Inversion of this ‘*perceptual model*’ produces subject-specific parameter estimates that determine the nature of the coupling between levels of the HGF. Inverting this model under generic (mean-field) approximations results in analytical belief-update equations, in which trial-wise belief updates are proportional to prediction errors (PEs) weighted by uncertainty (or its inverse, precision). The subject-specific parameters shape an individual’s approximation to ideal Bayesian inference, specifically how phasic and tonic volatility impacts trial-wise estimates of uncertainty at all levels of the hierarchy. Posterior estimates of HGF parameters can thus be regarded as a compact summary of an individual’s uncertainty processing during an experiment.

Furthermore, in a ‘*response model*’, trial-wise beliefs are probabilistically linked to observed trial-wise decisions. Inverting both perceptual and response models allows for estimating the parameters; this corresponds to Bayesian inference (of an observer) on Bayesian inference (of an agent) (Daunizeau *et al.*, 2010).

#### 3.5.2 The Perceptual Model

The goal of the HGF is to infer how an individual subject learns about hierarchically coupled environmental quantities under different forms of uncertainty (including volatility). In our case, the first quantity, *x*_1_, represents trial-wise outcomes (wins or losses) in the slot machine. This derives, through a sigmoid transform, from a second-level variable, *x*_2_ which represents, in logit space, the probability of winning (an indication of the slot-machine being ‘*hot*’ or ‘*cold*’ and likely (or not) to pay out). The variable *x*_2_ performs a Gaussian random walk trial by trial, with its step size (or variance) coupled to a higher level *x*_3_ (the speed at which a machine fluctuates between ‘*hot*’ and ‘*cold*’ states), according to f_2_(*x*_3_). The coupling between levels follows an expansion of log *f*(*x*) to first order, with subject-specific parameters *κ, ω* that determine an individual’s approximation to ideal Bayesian inference (see Equation 5 below). Finally, the parameter *ϑ* at the highest level denotes how quickly volatility itself is changing (meta-volatility). Generated observations, *u*, deterministically depend on *x*1, i.e., there is no sensory noise. A detailed derivation of the exact equations can be found in Mathys *et al.* (2014).

For model inversion and estimating posterior distributions of states and parameters, the HGF employs a generic variational Bayesian approximation, assuming that the posterior distributions of the states are Gaussian at all levels of the hierarchy:

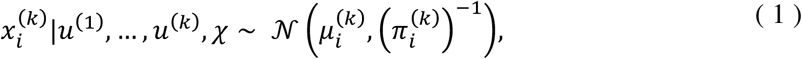

where u is an observed input, 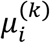 is the mean at time point *k* for level *i*, and 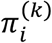 is the precision, or inverse variance, of this distribution, and 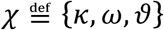 are subject-specific parameters. Updates to the posterior mean 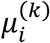 for level *i* have the general form:

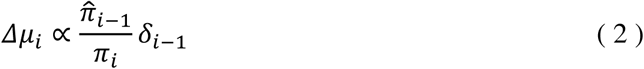

Thus, updates to the posterior mean at each level of the HGF are proportional to the prediction error (PE) at the level below, *δ*_*i*−1_, weighted by a ratio of uncertainties (or their inverses, precisions). Specifically, this ratio consists of the precision of the prediction onto the level below, 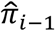, and the posterior precision at the current level, 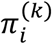.This equation reveals that the precision ratio corresponds to a dynamic learning rate: the higher the precision (the lower the uncertainty) of a prediction on the level below, the more meaningful the input from the level below and the greater the impact of the PE on updating the posterior mean. Conversely, the more certain an agent is about the true value of *x*_3_, the smaller the impact of the PE.

At the bottom of the hierarchy, the prediction error 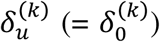 represents a value PE, the difference between the actual input *u* and the predicted input:

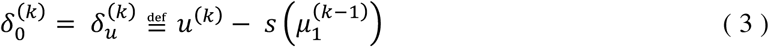

where *s* denotes the sigmoidal transform. At higher levels of the HGF, the PEs refer to volatility rather than value. A volatility PE (VOPE) integrates both predicted and observed, as well as informational and environmental uncertainty,

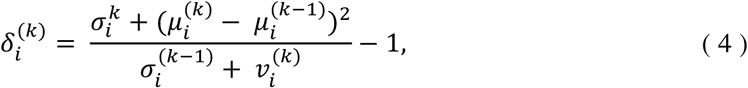

where 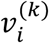 is defined as,

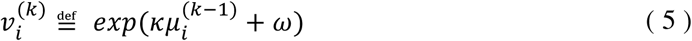

Here, informational uncertainty is given by 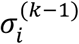, the variance of the posterior at trial *k*-1. Predicted total uncertainty is given by the denominator and includes both informational and environmental components. The environmental uncertainty represented by 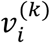 can be separated into phasic 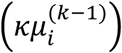 and tonic (*ω*) components, corresponding to ‘*unexpected*’ uncertainty related to environmental fluctuations. Additionally, *v* also affects the precision (or uncertainty) ratio (Equation 2) that defines the learning rate (for details, see Mathys *et al.* (2014)). Because the HGF can react sensitively to sudden shifts in environmental contingencies by adjusting the phasic component of its volatility estimate, it allows learning rates to be adjusted to the time-varying statistical structure of the environment. Individual differences in this dynamic learning process are determined by the agent-specific uncertainty parameters *κ, ω* and *ϑ*.

More concretely, in the context of our study, parameters *ω* and *ϑ* at the second and third level of the hierarchy, respectively, encode different aspects of subjective estimates of uncertainty. Specifically, these estimates concern environmental uncertainty, i.e., hidden fluctuations (volatility) of environmental states (Mathys *et al.*, 2014). These volatility estimates are potentially important for explaining the observed behaviour because they shape participants’ belief updates about the slot machine and their ensuing choices about gambling. Parameter *ω* represents a participant’s estimate of tonic volatility, i.e., how quickly a slot machine could be moving from a state where it is likely to pay out (running ‘*hot*’) to a state where it is not (running ‘*cold*’) and vice versa. Parameter *ϑ* encodes a participant’s estimate of meta-volatility, i.e., the tendency of volatility itself to change over time. Larger values of each parameter correspond to greater uncertainty in the participant’s perceptual inference process.

#### 3.5.3 The Response Model

The response model maps a subject’s beliefs (obtained by inverting the perceptual model under given parameter values) to observed gambling behaviour. Here, we use a sigmoidal response model (Mathys *et al.*, 2014). If this function is steep, there is a close relationship between current perceptual beliefs and betting behaviour. Conversely, a gentler sigmoidal slope results in a more stochastic mapping of beliefs to behaviour. This response function has a parameter, *β*, the decision ‘*temperature*’ (also known as the inverse temperature), that determines the steepness of the sigmoid and thus the degree of stochasticity in the belief-to-choice mapping. The larger the value of *β*, the steeper the function, and the more deterministic is the relationship between a subject’s belief and their actions. In this paper, we test the following two variants of this response model:

i. ‘*Standard*’ HGF: *β* = *constant*, i.e., the mapping from beliefs to behaviour is fixed across the experiment. This parameter is estimated for each participant.
ii. ‘*Uncertainty-driven*’ HGF: 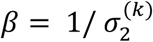, where 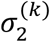 is the variance of the inferred probability of win on trial *k*. That is, the response behaviour dynamically adapts to the precision of the participants’ belief about the current probability of winning.

#### 3.5.4 Perceptual Variable

Based on previous work that examined different computational models of our slot machine paradigm (Paliwal et al., 2014), the perceptual variable used here was simple: a binary variable in which wins were represented by 1 and losses by 0 (Table 1). Although the task itself allows for continuous valued rewards, for this analysis, we look at wins and losses as binary outcomes (big wins and small wins are collapsed for the purpose of model fitting). We do so for the following reasons: first, we do not measure reward sensitivity on an individual basis, therefore creating a parametric multimodal reward variable that is consistent across the entire population is difficult. Secondly, measuring uncertainty-updating and impulsive responses in reaction to a binary win/loss event allows for a clearer interpretation of the model parameters and the correlational results.

**Table 1.** Composition of perceptual and response models for modelling.

| Composition of model variables | Win | Near-miss | True loss | BI | DU | CS | MS |
| --- | --- | --- | --- | --- | --- | --- | --- |
| Perceptual variables | 1 | 0 | 0 | - | - | - | - |
| Response Variables | - | - | - | 1 | 1 | 1 | 1 |

Variables are binary and composed on a trial-by-trial basis for each of combinations shown. BI = Bet increase; DU = double-up; CS = casino switch; MS = machine switch. Wins are encoded as 1, losses as 0.

#### 3.5.5 Response Variable

The response variable used in the model is constructed from four key behaviours afforded to participants as they play the virtual casino: bet behaviour, machine switching, doubling up and cashing out. We identify four key indicators of risk-taking within these behaviours:

i. Bet Increase (BI)
  – 1 = switching from a low to a high bet
  – 0 = staying at the same bet size
ii. Double Up (DU)
  – 1 = engaging in the secondary double-or-nothing option
  – 0 = declining to engage in the secondary double-or-nothing option
iii. Casino Switch (CS)
  – 1 = switching casino days
  – 0 = remaining in the current casino day
iv. Machine switch (MS)
  – 1 = deciding to switch to a new machine
  – 0 = continuing to play on the same machine

While these actions might at first glance appear to relate to different behaviours, they all share a common theme in that they enhance outcome variance and thus risk (compare the definition of risk in behavioural economics). For example, for a machine switch, regardless of whether the player is performing well or poorly on the current machine, the decision to switch machines incurs the risk that the new machine chosen may be punishing or rewarding, thereby making the player vulnerable to the variance of the task. Similarly, a bet increase is a risk-inducing shift in the face of uncertainty, again making the player more susceptible to larger wins and losses. And in the same respect, casino switches and double-ups again expose players to the risk that their environment will change dramatically, and for the worst. Each of the above actions thus leads to greater outcome variance (risk), and risk-taking, in turn, is one critical component of impulsivity (Whiteside and Lynam, 2001). In order to combine these behaviours into a single representation of risk taking, we construct a trial-wise, binary response variable per participant by performing an OR operation over the four actions. If a subject performs one or more of these actions on a given trial, the response variable on that trial is a 1. If the subject did not engage in any of these actions on a given trial, the response variable on that trial is a 0.

#### 3.5.6 Reinforcement Learning

As an alternative model, we used a classical associative learning model, Rescorla-Wagner (RW), often used in reinforcement learning (RL) (Rescorla and Wagner, 1972). The RW model updates the probability of a win on trial *k* by combining the probability on trial *k*-1 with a PE weighted by a constant learning-rate:

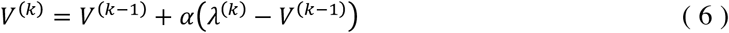

where *V*^(*k*)^ is the state of the tracked variable, in our case, the win probability, at time *k, V* ^(*k*−*1)*^ is the state of the variable at time point *k-1, α* is the learning rate and *λ*^(*k*)^ is the actual outcome at time point *k.*

Hence, in contrast to the HGF, the RW model does not have a dynamic learning rate over trials, nor can it account for different forms of perceptual uncertainty. Essentially, the RW model corresponds to an HGF with a fixed learning rate. Here, we combine the RW learning rule with the same sigmoidal response model described above, with free parameter *β*, that we estimate on a subject-specific basis. This results in a model that is (i) structurally not dissimilar but less complex than the HGF and (ii) almost identical to the RL model used in a prior investigation of learning after STN-DBS (Seymour *et al.*, 2016).

#### 3.5.7 Model Inversion

The HGF and RW models were inverted using population Markov-Chain Monte Carlo (MCMC) sampling (Aponte *et al.*, 2016). Parameter estimation in the HGF is classically ‘*fully Bayesian*’ and requires a selection of priors, which influence parameter estimation to a lesser or greater degree. In order to minimise this influence, we used a novel empirical Bayesian inference scheme for the HGF where a Gaussian group-level distribution of parameters is constructed from samples across the group. This group-level empirical prior is then used to obtain posterior parameter estimates in each subject (Figure 4). Participant-specific point estimates for model parameters are calculated as the median value of the participant’s posterior distribution.

**Figure 4.**
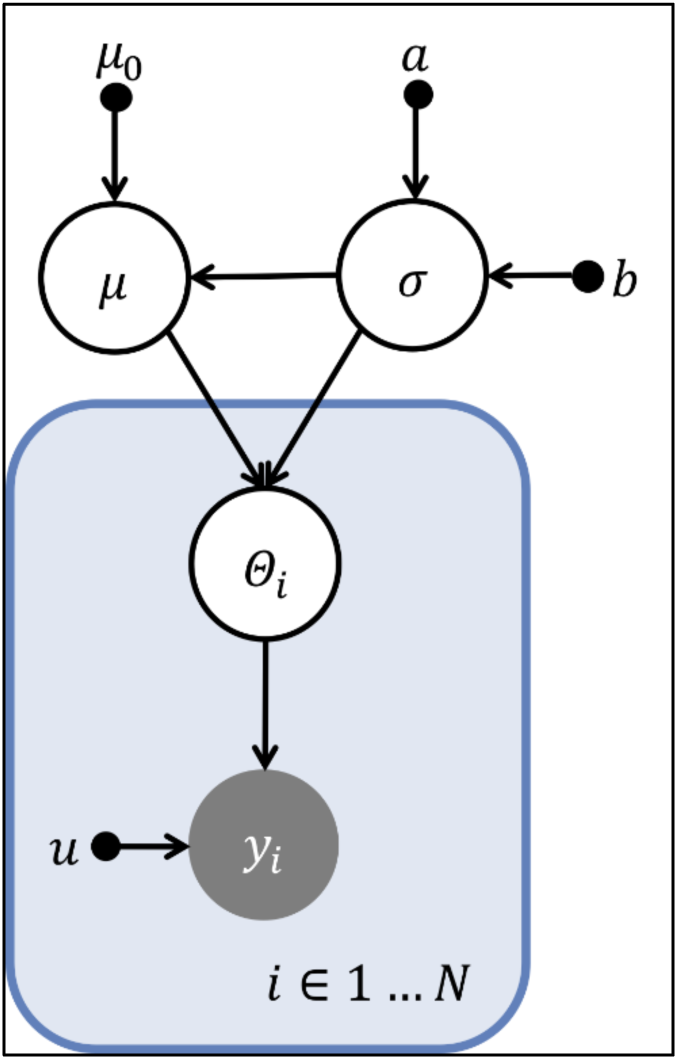
Schematic of hierarchical model inversion using population Markov-chain Monte Carlo sampling. Here, u represents the fixed perceptual variable that a participant observes during game play, and y_i_ represents the response variable for participant *i. Θ*_3_ = {*k, ω, ϑ, β*} is a participant-specific parameter vector consisting of the perceptual parameters shown in Figure 3 and the decision temperature, *β. μ* and *σ* are the mean and variance of the group-level empirical prior that is constructed using observations from all *N* participants. This prior is simultaneously used to invert models on a participant-specific level, for example, using u and y_1_, as the perceptual and response variables for Participant 1. In the notation used in this schematic, points indicate fixed parameters such as *b* or fixed data vectors such as *u*. Filled circles represent observed quantities, such as *y*_3_. Unfilled circles represent random variables that are estimated during model inversion. The hyperparameter for *μ, μ*_0_, is fixed to −3 for log *ω* and −6 for log *ϑ*, before inversion. The hyperprior on *σ* is a gamma distribution with parameters *a* and *b*, which are also fixed before inversion. As in previous work (Paliwal *et al.*, 2014), we did not estimate *κ* due to conditional dependencies with other parameters, but set it to unity.

Given the clinical constraints of our investigation (to reduce any burden on the participants, we only used 100 trials per episode of gambling, i.e., only half as many as in previous work) (Paliwal *et al.*, 2014), it was important to ensure that parameter estimates were robust. Therefore, in order to verify that HGF parameter estimates reliably reflected participant-specific characteristics of uncertainty-encoding and decision noise, we tested our ability to recover ground-truth parameter values from simulated response data. In order to assess parameter recoverability, we used three parameter values per parameter and generated a batch of thirty-eight synthetic response variables based on these assigned values, using the underlying trace of the slot machine as the perceptual variable. We then inverted the HGF and explored the relationship of the recovered parameter estimates, using the median of the posterior, with the ground truth values. When estimating *ω* and *ϑ, β* was held fixed; conversely, when estimating for *β, ω* and *ϑ* were fixed. This process was repeated for ten batches across each parameter.

#### 3.5.8 Model Comparison

As described above, we considered two competing hypotheses of how participants might incorporate uncertainty into their choice of actions, i.e., two different belief-to-choice mappings in the response model for the HGF (the ‘*Standard*’ and ‘*Uncertainty-driven*’ models). These two versions of the HGF were compared with the RW model. As we were primarily interested in the pre-DBS to post-DBS change, we selected the winning model for the pre-DBS measurements. We then evaluated if the parameter estimates of that winning model changed postoperatively. Estimates of the negative free energy (log model evidence) were computed using thermodynamic integration (Aponte *et al.*, 2016). The negative free energy balances goodness of fit with a complexity penalty. Group-level free energy estimates were compared to select a winning model.

### 3.6 Data Analysis

#### 3.6.1 General Considerations

All computational modelling and model inversion was performed using MATLAB (Mathworks), employing custom scripts developed from the HGF toolbox (http://www.translationalneuromodeling.org/tapas/). Multiple regression analyses were performed using the regstats function in the MATLAB Statistics Toolbox. For all analyses involving multiple comparisons, native *p*-values are presented, accompanied by Holm-Bonferroni correction at *α*=0.05. To test the significance of individual regressors in multiple regression models, post hoc t-tests were performed.

Neuropsychiatric assessment data from baseline, prior to DBS, was compared with data gathered at thirteen-weeks post-DBS, when the gambling task was repeated. To test for differences in pre-DBS and post-DBS questionnaire scores and model parameter estimates, a paired t-test was employed when the data were normally distributed and the Wilcoxon signed-rank test otherwise, where distribution was assessed using the Lilliefors test. Gambling behaviours (such as bet increases and machine switches) were also compared at both intervals. Gambling behaviours were regressed against clinical measures of impulsivity to determine significant relations. After determining the winning computational model, model parameter estimates were extracted for each participant and regressed against clinical measures of impulsivity to determine significant associations and predictors of postoperative impulsivity. Based on previous work showing a significant association between BIS scores and both slot machine behaviour and HGF-based estimates of uncertainty encoding (Paliwal *et al.*, 2014), we focused our analyses on the BIS and its subscales. Perceptual model parameters were extracted in log space: *ω* and *β* are naturally estimated in log space, since they are part of exponential terms in their respective equations (see equation 5).

From a clinical perspective, we were interested in examining whether changes in the computational characterisation of individual uncertainty estimates pre- to post-DBS were associated with clinically-relevant changes in impulsivity at any time point after DBS. Our strategy to attempt prediction of clinical outcomes follows the ‘*generative embedding*’ approach, in which individual predictions are not derived from measured data but from parameter estimates obtained by a generative model (Brodersen *et al.*, 2011; Brodersen *et al.*, 2014). Importantly, stimulation-dependent changes in impulsivity may evolve in an unpredictable manner subsequent to DBS, related to variations in DBS programming over time (with considerable adjustments to stimulation in the first six postoperative months). Furthermore, the optimal BIS cut-off score for clinically-significant impulsivity varies by age and disease (Stanford *et al.*, 2009), with only one existing investigation specific to a Parkinson’s disease cohort (Voon *et al.*, 2007). Therefore, we examined whether individual changes in parameter estimates associated with the maximum postoperative increase in impulsivity, as measured by the BIS, compared to baseline, across six months of longitudinal follow up.

## 4 Results

### 4.1 Participant Characteristics

Participants were a predominantly middle-aged sample, with a bias towards male gender and akinetic-rigid/mixed phenotype over tremor (Table 2). Most participants had bilateral disease with consequent impairment of functioning in their activities of daily living.

**Table 2.** Demographic and clinical characteristics of cohort.

| **Categorical Variable** | **Total (n=38)** |  |
| --- | --- | --- |
| **Gender** | ***n*** | **% total** |
| Male | 25 | 65.8 |
| Female | 13 | 34.2 |
| **Clinical Subtype** | ***n*** | **% total** |
| Akinetic-Rigid | 13 | 34.2 |
| Mixed | 18 | 47.4 |
| Tremor | 7 | 18.4 |
| **Continuous Variable** | ***Mean (SD), Median (Range)*** |  |
| **Age (Years)** | 61.9 ($\pm 9.3$),<br>65 (35 - 76) | |
| **Hoehn & Yahr Stage** | 2.7 ($\pm 0.6$),<br>2.5 (1.5 - 4) | |
| **Years Since Diagnosis** | 8.5 ($\pm 4.6$),<br>7 (2 - 21) | |

### 4.2 Neuropsychiatric Assessment Pre- and Post-DBS

Concerning symptoms of primary interest (Table 3), there was a small but statistically-significant group-level post-DBS decrease in impulsivity, as measured by the BIS Total, compared to baseline. There was also a significant reduction of motor symptoms assessed using the UPDRS Part III Motor Examination, with a corresponding significant reduction in the requirement for dopaminergic therapy (LEDD). There were no statistically-significant changes in other behavioural measures related to impulsivity, including the Hayling test, the Excluded Letter Fluency task and the delay discounting task. Comparable to the BIS, the QUIP-RS total score demonstrated a trend towards a reduction at thirteen-weeks post-DBS, but this did not reach significance.

**Table 3.** Neuropsychiatric assessment data pre- and post-DBS. ‡Indicates significant native *p*-values, before multiple comparison correction Significance: *** p<0.001, ** p<0.01, * p<0.05, where *p*-values are Holm-Bonferroni corrected for multiple comparisons with *α* = 0.05.

| Behavioural Measure | Pre-DBS | Post-DBS | Max Impairment | Pre- vs. Post-DBS |  |  |
| --- | --- | --- | --- | --- | --- | --- |
| Gaussian Distribution | Mean (SD), Median (Range) | | | $t$-stat | $p$-value | Adj. $p$-value |
| BIS | 60.2 ($\pm 7.4$),<br>60 (44 - 76) | 57.8 ($\pm 9.5$),<br>58 (40 - 75) | 2.3 ($\pm 6.4$),<br>3 (-14 - 17) | 2.66 | 0.011 ‡ | 0.033* |
| BDI | 11.1 ($\pm 5.1$)<br>11 (1 - 22) | 8.4 ($\pm 6.6$)<br>6 (1 - 29) | 1.3 ($\pm 7.1$),<br>0 (-12 - 18) | 2.29 | 0.028 ‡ | 0.056 |
| QUIP-RS | 21.4 ($\pm 15.7$),<br>19 (0 - 63) | 17.2 ($\pm 15.8$),<br>14 (0 - 55) | 5.1 ($\pm 13.5$),<br>4 (-25 - 37) | 1.78 | 0.084 | 0.084 |
| UPDRS Part III Motor | 37.7 ($\pm 17.0$),<br>37 (10 - 91) | 30.9 ($\pm 12.7$),<br>32 (8 - 60) | N/A | 3.47 | 0.001 ‡ | 0.004** |
| LEDD | 1032.9 ($\pm 599.4$),<br>988 (0 - 3450) | 334.9 ($\pm 199.8$), 329<br>(0 - 825) | N/A | 8.59 | <0.001 ‡ | <0.001*** |
| Skewed Distribution | Mean (SD), Median (Range) | | | Chi-square | $p$-value | Adj. $p$-value |
| ELF Rule Violations | 9.7 ($\pm 5.3$),<br>9 (0 - 24) | 8.4 ($\pm 5.0$),<br>8 (1 - 18) | 3.0 ($\pm 6.5$),<br>1 (-6 - 19) | 20.2 | 0.12 | 0.360 |
| Hayling AB Error Score | 11.4 ($\pm 11.1$),<br>8 (0 - 38) | 9.4 ($\pm 11.3$),<br>5 (0 - 45) | 6.2 ($\pm 12.0$),<br>6 (-19 - 30) | 23.6 | 0.13 | 0.360 |
| Delay Discount K | 0.034 ($\pm 0.067$),<br>0.016 (0.00016 -<br>0.25) | 0.036 ($\pm 0.049$),<br>0.016 (0.00016 -<br>0.25) | 0.041 ($\pm 0.077$),<br>0.0128 (-0.15 - 0.25) | 12.6 | 0.18 | 0.260 |

Abbreviations: BDI = Beck Depression Inventory, BIS = Barratt Impulsiveness Scale, ELF = Excluded Letter Fluency, LEDD = Levodopa Equivalent Daily Dose, QUIP-RS = Questionnaire for Impulsive-Compulsive Disorders in Parkinson's Disease Rating Scale, UPDRS = Unified Parkinson's Disease Rating Scale
Max impairment refers to the maximum impairment for each outcome across all measurement intervals subsequent to DBS.

For symptoms of secondary interest (and subscales), see Supplementary Table 1. There was considerable variance between subjects across assessment scores at each interval and within subjects across the course of longitudinal follow up (Supplementary Figure 1).

The BIS and the BDI showed a significant positive correlation at each time point (*ρ*_*pre*_=0.46, *p*=0.003; *ρ*_*post*_ =0.53, *p*<0.001), and both showed (near-)significant changes from pre- to post-DBS (Table 8.3). Therefore, to rule out that impulsivity-related findings were driven by changes in depression, the BDI was included as a covariate when regressing behaviour and model parameter estimates against BIS scores. Whilst the LEDD is conceivably related to impulsivity, it did not correlate with the BIS total (*ρ*_*PRE*_=-0.126, *p*=0.450; *ρ*_*post*_=-0.042, *p*=0.799) and was therefore not included in these regression analyses. However, the QUIP and LEDD correlated strongly at both time points (*ρ*_*PRE*_=0.42, *p*=0.008); *ρ*_*post*_ =0.44, *p*=0.005), with LEDD decreasing significantly post-DBS. There were no significant correlations between LEDD and the other measures of impulsivity (ELF Rule Violations, Hayling AB Error Score and Delay Discount K). Based on previous work using this task and modelling framework (Paliwal *et al.*, 2014), we focused our attention on exploring impulsivity as measured by the BIS.

### 4.3 Gambling Behaviour

#### 4.3.1 Gambling Behaviour Pre- and Post-DBS

At the group level, there were no significant differences in the behaviour of participants on the slot machine from pre- to post-DBS (Table 4). Due to subjects not engaging in the casino switch option, this variable was eliminated from regression analyses.

**Table 4.** Gambling behaviour, pre- and post-DBS.

| Gambling Behaviour | Pre-DBS | Post-DBS | Pre- vs. Post-DBS |
| --- | --- | --- | --- |
|  | Mean (SD, Range) |  | t-stat (p-value) |
| Bet Increases | 39.732 ($\pm 42.98$; 5-174.4) | 54.896 ($\pm 69.261$; 5-339.75) | -1.873 (0.069) |
| Machine Switches | 2.052 ($\pm 2.968$; 0-12) | 1.079 ($\pm 1.978$; 0-10) | 1.837 (0.074) |
| Cashout | 0.136 ($\pm 0.342$; 0-1) | 0 ($\pm 0$; 0-0) | N/A |
| Double-up | 10.736 ($\pm 9.841$; 0-25) | 13.132 ($\pm 9.510$; 0-25) | -1.483 (0.147) |

#### 4.3.2 Pre-DBS Regression of BIS scores on Gambling Behaviour

We studied the relationship between pre-DBS gambling behaviour and pre-DBS impulsivity as measured by the BIS (Table 5). The BDI was included in this regression in order to control for changes in clinical state attributable to depressive symptoms. The overall preoperative model including the BDI total was significantly associated with the BIS total score [*F*_(4,33)_=3.024, *p*=0.031]. Post-hoc t-tests on task behaviour revealed that no behavioural variable was significantly related to the BIS individually. When subscales of the BIS were examined, gambling behaviour associated significantly with the BIS Attentional subscale [*F*_(4,33)_=4.094, *p*=0.008], where higher bet sizes corresponded to higher attentional impulsivity (*t*_(37)_=2.303, *p*=0.028) (Supplementary Table 2).

**Table 5.** Pre-DBS slot machine behaviour and pre-DBS BIS. *b* values are standardized regression coefficients. Significance: *** p<0.001, ** p<0.01, * p<0.05, where *p*-values are Holm-Bonferroni corrected for multiple comparisons with *α* = 0.05. ^ Indicates significant t-statistics, Holm-Bonferroni corrected for multiple comparisons.

|  | Independent Variables (*b*) |  |  |  |  |  |  |
| --- | --- | --- | --- | --- | --- | --- | --- |
| Dependent Variable | Bet Size | Machine Switch | Double-up | BDI | $R^2$ | F-stat | *p*-value |
| BIS Total | 0.034 | 0.338 | -0.077 | 0.821 | 0.268 | 3.024 | 0.0341* |

#### 4.3.3 Post-DBS Regression of BIS scores on Gambling Behaviour

The full model of postoperative gambling behaviour was also significantly associated with BIS total score [*F*_(4,33)_=4.920, *p*=0.003] (Table 6). Again, post-hoc t-tests revealed that no task behaviour was significant on its own. When subscales of the BIS were examined, gambling behaviour correlated significantly with the BIS Attentional subscale [*F*_(4,33)_=8.123, *p*<0.001]. Post-hoc t-tests revealed that higher bet sizes (*t*_(37)_=2.604 *p*=0.014) and more frequent double or nothing gambles (*t*_(37)_=2.589 *p*=0.014) corresponded to higher BIS Attentional scores (Supplementary Table 3).

**Table 6.** Post-DBS slot machine behaviour and post-DBS BIS. *b* values are standardized regression coefficients. Significance: *** p<0.001, ** p<0.01, * p<0.05, where *p*-values are Holm-Bonferroni corrected for multiple comparisons with *α* = 0.05. ^ Indicates significant t-statistics, Holm-Bonferroni corrected for multiple comparisons.

|  | Independent Variables (*b*) |  |  |  |  |  |  |
| --- | --- | --- | --- | --- | --- | --- | --- |
| Dependent Variable | Bet Size | Machine Switch | Double-up | BDI | $R^2$ | F-stat | *p*-value |
| BIS Total | 0.018 | 0.128 | 0.263 | 0.712 | 0.374 | 4.920 | 0.003** |

Post-DBS, higher bets and more frequent machine switches were significantly associated with higher QUIP-RS scores (Supplementary Table 4). No other measures of impulsivity were significantly associated with pre- or post-DBS slot machine activity.

#### 4.3.4 Maximum BIS Increase and Perioperative Changes in Gambling Behaviour

The change in gambling behaviours between the pre- and post-DBS time points were significantly associated with the maximum postoperative increase in BIS score. [*F*_(4,33)_=3.516, *p*=0.017] (Table 7). Post-hoc t-tests revealed that the change in bet behaviour significantly was associated with maximum BIS increase (*t*_(37)_=2.866, *p*=0.007). Additionally, the change in machine switch behaviour was also significantly associated with maximum BIS increase (*t*_(4,33)_=2.219, *p*=0.034). In other words, changes in betting and slot machine switching behaviours after DBS indexed changes in impulsivity as assessed by the BIS.

**Table 7.**
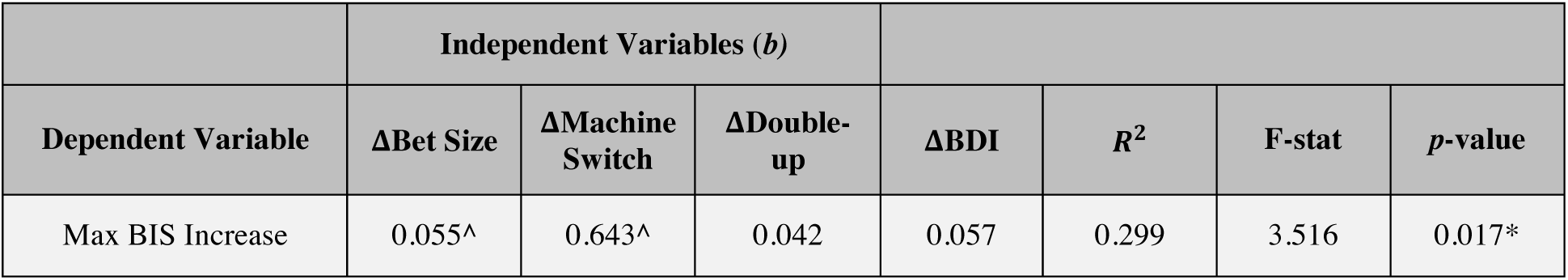
Pre-to-Post-DBS slot machine behaviour and max BIS increase. *b* values are standardized regression coefficients. Significance: *** p<0.001, ** p<0.01, * p<0.05, where *p*-values are Holm-Bonferroni corrected for multiple comparisons with *α* = 0.05. ^ Indicates significant t-statistics, Holm-Bonferroni corrected for multiple comparisons.

|  | Independent Variables (*b*) |  |  |  |  |  |  |
| --- | --- | --- | --- | --- | --- | --- | --- |
| Dependent Variable | ΔBet Size | ΔMachine Switch | ΔDouble-up | ΔBDI | $R^2$ | F-stat | *p*-value |
| Max BIS Increase | 0.055^ | 0.643^ | 0.042 | 0.057 | 0.299 | 3.516 | 0.017* |

### 4.4 Computational Modelling

As described above, we were interested in evaluating the role of uncertainty and its association with postoperative changes in impulsivity in our cohort. We therefore first determined, using Bayesian model comparison, which of our three models best explained pre-DBS behaviour, before evaluating whether the parameter estimates of this winning model changed postoperatively and were associated with postoperative BIS scores. Bayesian model comparison selected the ‘*standard*’ HGF (with a participant-specific decision temperature in the response model) as the winning model, with a group-level Bayes factor of approximately 12.5, compared to the next best model (the Rescorla-Wagner model) (Figure 5).

**Figure 5.**
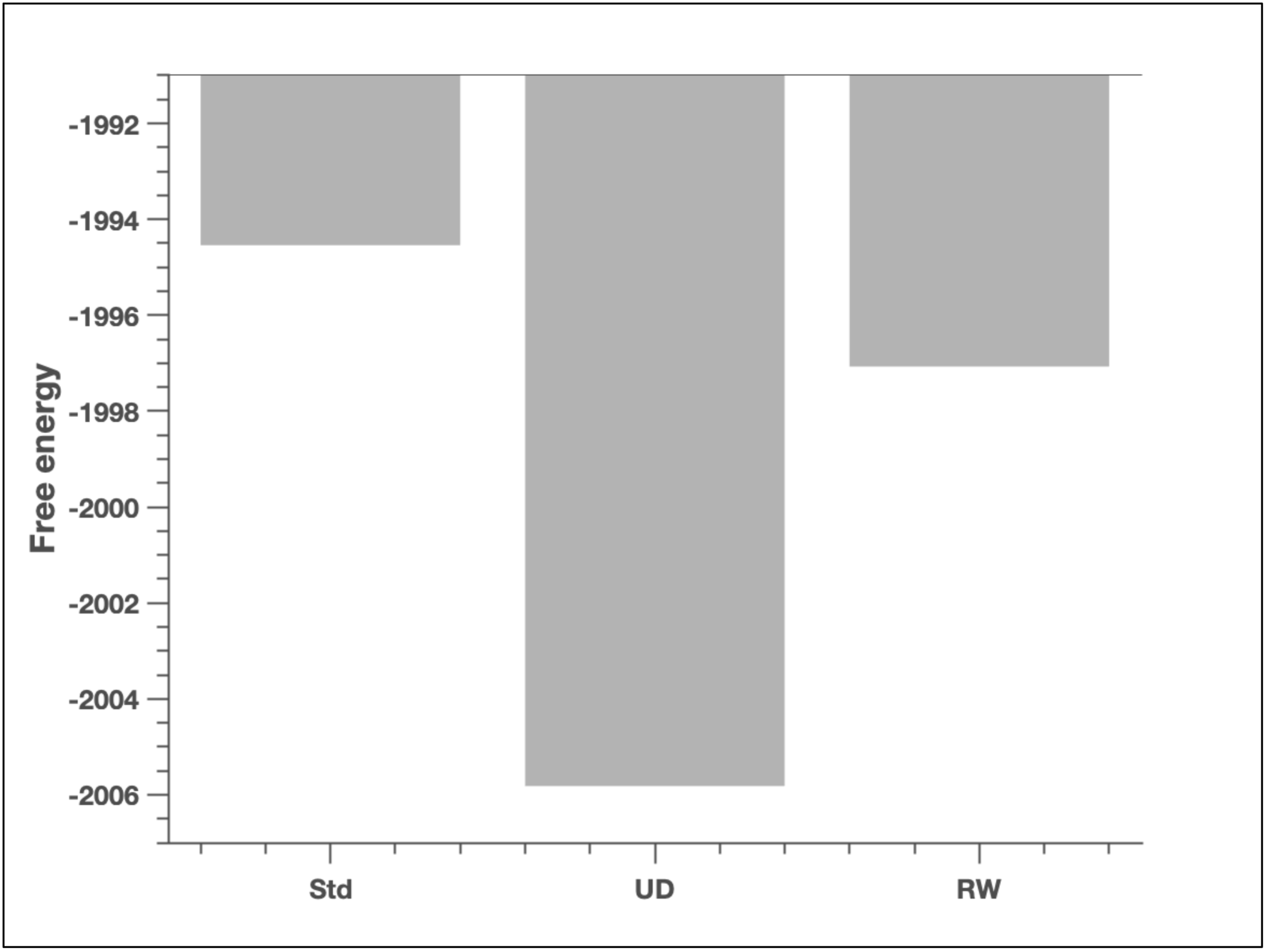
Model comparison results. Bayesian model comparison results across the Standard (Std), Uncertainty-driven (UD) and Rescorla Wagner (RW) models, pre-and post-DBS. Shown here are the group-level free energy values for the three models, Std, UD and RW. Pre-DBS model free energies are *F*_*Std*_ =-1994.54, *F*_*UD*_=-2005.82, and *F*_*RW*_=-1997.07. The winning model pre-DBS is the standard HGF. The group-level difference in free energy compared to the next best model (the Rescorla-Wagner model) is 2.53, corresponding to a Bayes factor of approximately 12.5.

#### 4.4.1 Parameter recoverability in the HGF

We tested for parameter recoverability in the HGF, finding that ground truth parameter values for *ω* and *β* could be recovered consistently, but we were unable to reliably recover *ϑ* (Figure 6). For this reason, we restricted the following analysis to parameters *ω* and *β* when exploring the association between parameter values and questionnaire-based measures of impulsivity.

**Figure 6.**
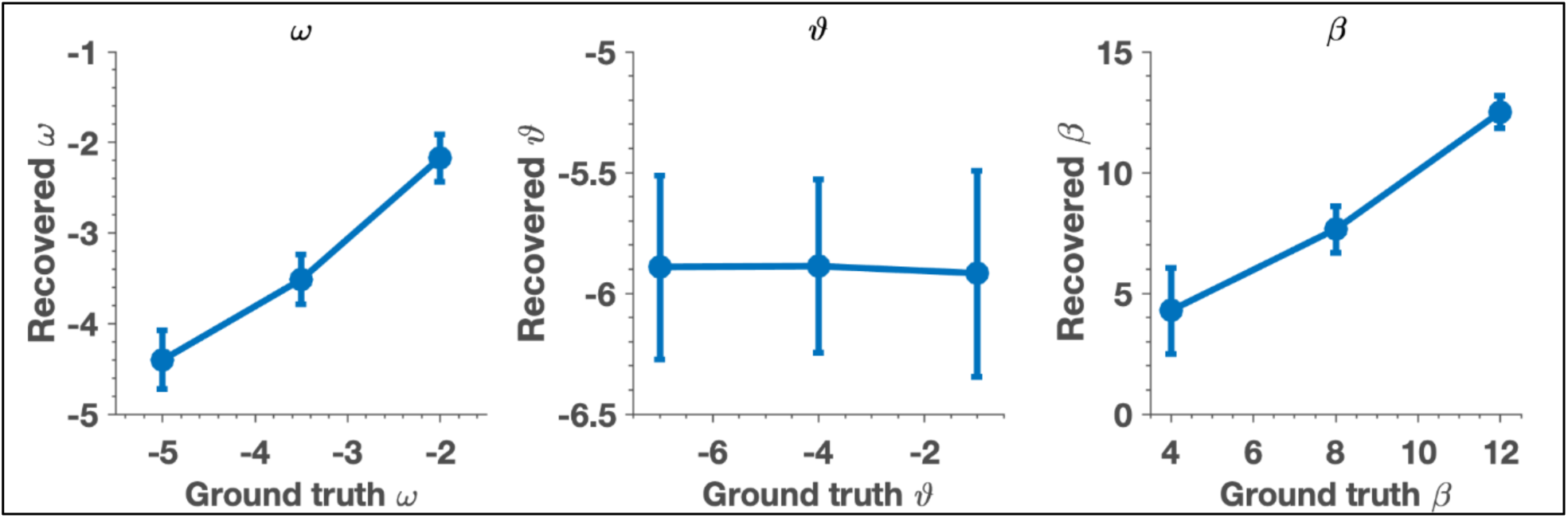
Parameter recoverability in the HGF. The first plot above shows the mean of the recovered parameter values for *ω* across 10 inversions of the HGF on the y-axis, against the ground truth *ω* values on the x-axis. The second plot shows the mean of recovered parameter values for *ϑ* across 10 inversions of the HGF on the y-axis, against the ground truth *ϑ* values on the x-axis. The third plot shows the mean of the recovered parameter values for *β* across 10 inversions of the HGF on the y-axis, against the ground truth parameters for *β* on the x-axis. Error bars indicate the standard deviation of the recovered parameter estimates across 10 inversions.

#### 4.4.2 Changes in Model Parameter Estimates Pre- to Post-DBS

Estimates of the HGF model perceptual parameter *ω* significantly increased postoperatively (*t*_37_ =-61.328, p<0.001), and estimates of *β* significantly decreased (*t*_37_=2.124, p=0.04), implying larger subjective estimates of uncertainty (volatility) and greater stochasticity in the selection of responses after DBS (Table 8 and Figure 7). *ω* represents a subject’s estimate about the tonic component of environmental volatility; i.e., how quickly the likelihood of winning on a given slot machine might be changing, while *β* represents the decision noise, or the stochasticity involved in the belief-to-choice mapping process.

**Table 8.** HGF model parameters, pre- and post-DBS. Group means are reported with standard deviations and range in parentheses. Model parameters are reported in log space. Significance: *** p<0.001, ** p<0.01, * p<0.05, where p-values are Holm-Bonferroni corrected for multiple comparisons with *α* = 0.05.

| Parameter | Pre-DBS | Post-DBS | t-stat | p-value |
| --- | --- | --- | --- | --- |
| $\omega$ | -8.165 (0.207, -8.740--7.843) | -5.491 (0.261, -6.187--4.952) | -61.328 | <0.001*** |
| $\beta$ | 3.279 (2.802, 0.160-10.157) | 2.218 (2.525, 0.096-9.721) | 2.124 | 0.04* |

**Figure 7.**
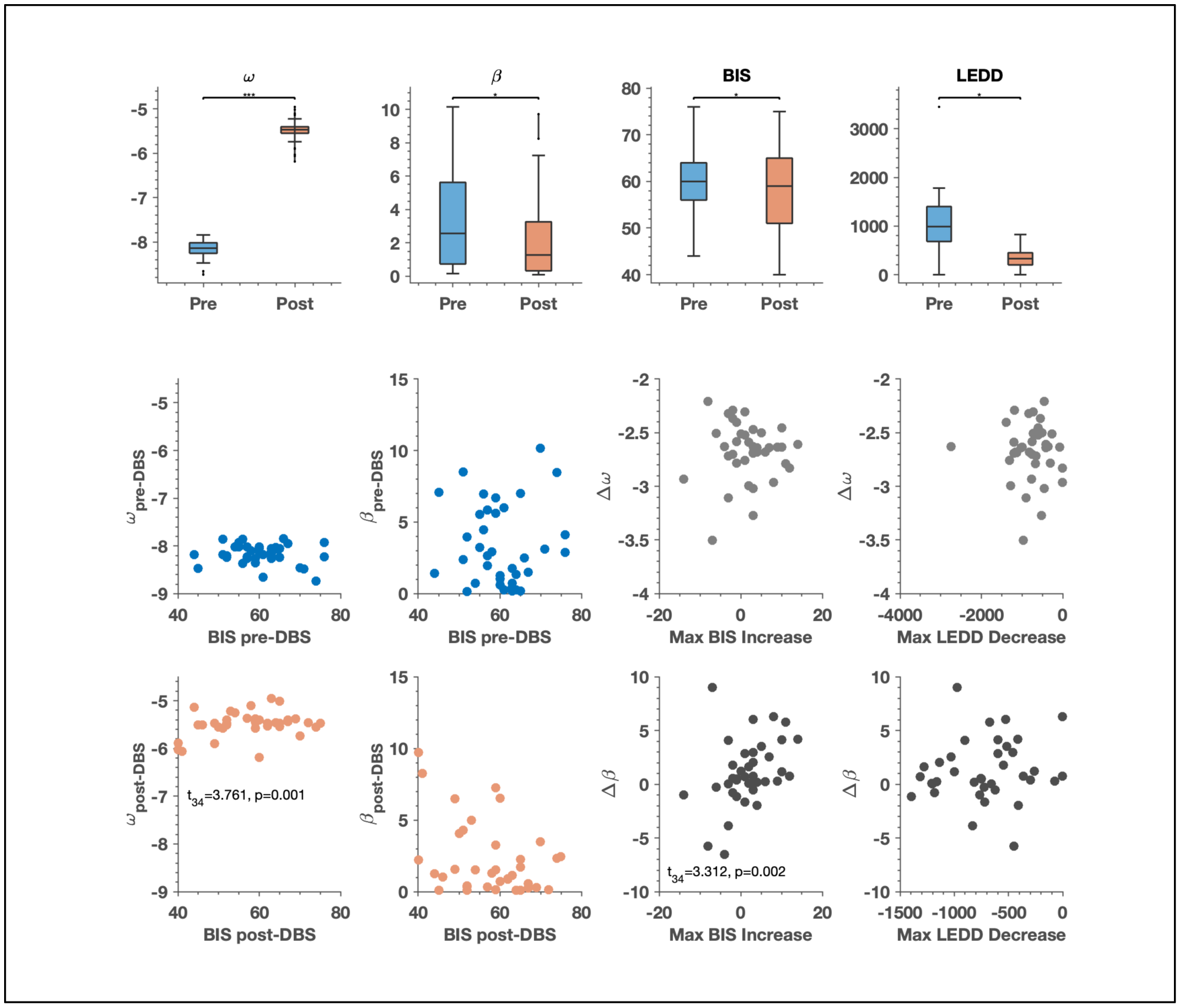
Computational model parameters, pre- and post-DBS. In the first row of figures, model parameter estimates for *ω* and *β*, BIS scores and LEDD scores are displayed pre- and post-DBS. For the box plots, the central line indicates the median of the distribution, and the top and bottom edges of the box represent the 25^th^ and 75^th^ percentiles respectively. The whiskers extend to the farthest data points that are included in the distribution and are not considered outliers. A paired t-test was performed to determine the pre-DBS vs. post-DBS difference in the distributions of *ω, β*, the BIS and the LEDD. Significant differences were observed in estimates of parameter *ω* (*t*_(37)_ =-61.328, *p*<0.001) and in estimates of parameter *β* (*t*_(37)_ =-2.214, *p*=0.04) as shown in Table 8. Also shown is the change in BIS pre- and post-DBS (*t*_(37)_ =-2.66, *p*=0.033), as shown in Table 3. *p*-values are Holm-Bonferroni corrected for multiple comparisons with *α* = 0.05. The second row illustrates the relationship between pre-DBS *ω* and pre-DBS BIS, pre-DBS *β* and pre-DBS BIS and the pre-to-post change in *ω* with the max increase in BIS, as well as the max decrease in LEDD. The third row illustrates the relationship between post-DBS *ω* and post-DBS BIS, post-DBS *β* and post-DBS BIS and the pre-to-post change in *β* with the max increase in BIS. Here, we have removed the outlier in the plot relating the change in *β* to the max decrease in LEDD. These plots serve to better illustrate the results shown in Tables 10 and 11. Specifically, that greater volatility estimates (*ω*, the tendency of a slot machine’s winning probability to change) were associated with greater maximum postoperative BIS scores, and that greater stochasticity in belief-to-choice mapping (decision temperature - *β*) associated significantly with the maximum postoperative increase in BIS.

#### 4.4.3 Pre-DBS Regression of BIS scores on Model Parameter Estimates

The full regression model (including the estimates of preoperative perceptual and response parameters *ω* and *β*) was significantly associated with BIS total [*F*_(3,34)_=3.372, *p*=0.03] (Table 9). Post-hoc t-tests on model parameter estimates did not reveal any single parameter to be significantly related to the BIS on its own. When subscales of the BIS were examined, the full regression model associated significantly with the BIS Attentional subscale [*F*_(3,34)_=3.314, *p*=0.031], but again no single parameter was independently significant (Supplementary Table 5).

**Table 9.** Pre-DBS model parameters and pre-DBS BIS. *b* values are standardized regression coefficients. Significance: *** p<0.001, ** p<0.01, * p<0.05, where *p*-values are Holm-Bonferroni corrected for multiple comparisons with *α* = 0.05. ^ Indicates significant t-statistics, Holm-Bonferroni corrected for multiple comparisons.

| Dependent Variable | Independent Variables ($b$) | | | | | |
| --- | --- | --- | --- | --- | --- | --- |
| | $\omega$ | $\beta$ | BDI | $R^2$ | F-stat | p-value |
| BIS-11 Total | -5.306 | -0.370 | 0.729 | 0.229 | 3.372 | 0.03* |

**Table 10.** Post-DBS model parameters and post-DBS BIS. *b* values are standardized regression coefficients. Significance: *** p<0.001, ** p<0.01, * p<0.05, where *p*-values are Holm-Bonferroni corrected for multiple comparisons with *α* = 0.05. ^ Indicates significant t-statistics, Holm-Bonferroni corrected for multiple comparisons.

| Dependent Variable | Independent Variables (*b*) |  |  |  |  |  |
| --- | --- | --- | --- | --- | --- | --- |
| | $\omega$ | $\beta$ | BDI | $R^2$ | F-stat | *p*-value |
| BIS Total | 17.434^ | -0.007 | 0.943 | 0.49 | 10.906 | <0.001*** |

**Table 11.** Max BIS increase and change in model parameters. *b* values are standardized regression coefficients. Significance: *** p<0.001, ** p<0.01, * p<0.05, where *p*-values are Holm-Bonferroni corrected for multiple comparisons with *α* = 0.05. ^ Indicates significant t-statistics, Holm-Bonferroni corrected for multiple comparisons. *Changes in ω and β are calculated as the pre-DBS parameter estimate value minus the post-DBS parameter estimate value. Max BIS increase refers to the maximum impairment for impulsivity across all measurement intervals subsequent to DBS.*

|  | Independent Variables (*b*) |  |  |  |  |  |
| --- | --- | --- | --- | --- | --- | --- |
| Dependent Variable | $\Delta\omega$ | $\Delta\beta$ | $\Delta\text{BDI}$ | $R^2$ | F-stat | *p*-value |
| Max BIS Increase | 8.681 | 1.220^ | -0.098 | 0.260 | 3.987 | 0.015* |

#### 4.4.4 Post-DBS Regression of BIS scores on Model Parameter Estimates

The full regression model was significantly associated with BIS total [*F*_(3,34)_=10.906, *p*<0.001] (Table 10). Post-hoc t-tests revealed that *ω* was a significant regressor (*t*_(34)_=3.761, *p*=0.001). The positive regression coefficient for *ω* implies that the greater the subjective estimate of uncertainty (tonic volatility), the higher the BIS score (Figure 7). In other words, although there was a group level decrease in BIS score pre-to-post DBS, at an individual level: the greater postoperative estimate of uncertainty (i.e., the higher the estimated volatility of the slot machine’s winning probability), the greater the postoperative impulsivity. Thus, for participants with high subjective volatility estimates, there was likely to be a postoperative increase in BIS. When subscales of the BIS were examined, model parameter estimates correlated significantly with the BIS Attentional subscale [*F*_(3,34)_=7.777, *p*<0.001] and the BIS Non-Planning subscale [*F*_(3,34)_=7.642, *p*<0.001]. Post-hoc t-tests revealed that *ω* was also a significant regressor in both of these associations (*t*_(34)_=2.874, *p*=0.007 for BIS Attentional and *t*_(34)_=2.561, *p*=0.015 for BIS Non-Planning) (Supplementary Table 6).

#### 4.4.5 Change in Model Parameters Associating with Maximum Change in BIS

We were interested in whether individual pre- to postoperative changes in estimates of subjective uncertainty would correlate with the maximum postoperative change in impulsivity across six months of follow up post-DBS. The pre-to-post change in model parameter estimates was calculated as the preoperative minus the postoperative parameter estimate. In the case of *ω*, postoperative parameter estimates were significantly higher than preoperative values (in the case of *ω*, higher values imply higher uncertainty), therefore the Δ*ω* value for each subject is negative and implies a post-operative increase in uncertainty. With regards to *β*, postoperative parameter estimate values were significantly lower than preoperative values (in the case of *β*, lower values imply greater stochasticity), therefore the Δ*β* value for most subjects is positive, implying a postoperative increase in the stochasticity of the belief-to-choice mapping. Changes in model parameter estimates were associated with the maximum post-operative increase in the BIS across all longitudinal assessments over six months [*F*_(3,34)_= 3.987, *p*=0.015] (Table 11). Post-hoc t-tests revealed that Δ*β* was a significant regressor (*t*_(34)_=3.312, *p*=0.002). The positive regression coefficient here implies that a post-operative increase in decisional randomness related to a greater maximum increase in BIS (Figure 7).

#### 4.4.6 Supplementary Analyses

We examined whether dopaminergic medication dosage expressed as a standardised unit (LEDD) was connected to computational model parameters and whether changes in drug doses postoperatively were connected to changes in uncertainty encoding. There was no significant relationship between pre-DBS model parameters and pre-DBS LEDD [*F*_(2.35)_= 1.008, *p*=0.375] or post-DBS model parameters and post-DBS LEDD [*F*_(2.35)_= 0.266, *p*=0.768]. Additionally, the pre-to-postoperative change in LEDD did not significantly relate to pre-to-postoperative change in BIS (*ρ*=0.28, *p*=0.088). Also, the maximum decrease in LEDD across six months of follow up did not significantly relate to the postoperative maximum increase in BIS (*ρ*=0.29, *p*=0.073),

However, there was a significant relationship between the change in model parameter estimates and the maximum post-operative decrease in LEDD [*F*_(2.35)_=4.032, *p*=0.027] (Supplementary Table 7). The BDI was not included in these regression models, as it is not a confound when examining a relationship between model parameter estimates and the LEDD. Post-hoc t-tests showed that Δ*β* was a significant regressor (*t*_(34)_=2.832, *p*=0.008). The positive regression coefficient here implies that a post-operative increase in decisional stochasticity was observed in patients who had a larger post-operative decrease in LEDD. However, this relationship appeared to be driven by an outlier participant with a particularly large perioperative decrease in LEDD. When this participant was removed, the relationship was no longer statistically significant. We have removed the outlier in Figure 7 but a full plot including the outlier can be found in Supplementary Figure 2.

## 5 Discussion

In this study, we employed a naturalistic gambling task and a hierarchical Bayesian model (for inference on subject-specific estimates of uncertainty) in order to investigate impulsive decision-making in participants with Parkinson’s disease undertaking subthalamic DBS. Gambling behaviour associated with a ‘*gold-standard*’ questionnaire (BIS) measure of impulsivity, with post-DBS changes in gambling behaviours indexing postoperative changes in impulsivity. We also found that parameter estimates representing subjective estimates of environmental uncertainty (volatility) changed significantly from pre- to postoperative conditions. In particular, there was a significant increase in *ω*, that reflects a gambler’s estimate of how quickly the probability of winning on a given slot machine was changing (volatility), There was also a postoperative decrease in a second parameter, *β*, that captures the decision noise in a player’s belief-to-choice mapping. Notably, these model-based estimates of uncertainty related to postoperative impulsivity. The greater the postoperative estimate of *ω*, the greater the postoperative BIS score. In other words, the more a participant perceived the pay-out tendency of a slot machine to be changing after DBS, the more impulsive they rated themselves. Additionally, the higher the pre- to postoperative decrease in estimates of *β*, the higher the postoperative increase in BIS score across six months of longitudinal follow up. In other words, the more a participant became indiscriminate in their belief-to-choice mapping after DBS, the more impulsive they rated themselves.

Our gambling task utilised a multivariate response variable (bet increase, machine switch, casino switch and double-up) that captured different aspects of impulsivity and explorative behaviour. Furthermore, by employing a generative model that mapped observed responses to perceptual states, we were able to infer directly upon participant-specific parameters defining individual differences in uncertainty encoding. This is an important point of difference from a purely behavioural analysis, in which responses can have more than one (ambiguous) proximate cause. In the HGF, parameters are mathematically defined and have a concrete influence upon learning at different levels of the hierarchy (Mathys *et al.*, 2011; Mathys *et al.*, 2014).

What is the significance of individual differences in uncertainty encoding? Increased estimates of environmental uncertainty accelerate the rate of learning at higher hierarchical levels, which could engender maladaptive learning at lower levels of the hierarchy. A high learning rate suppresses the influence of top-down expectations, and may impair learning about probabilistically aberrant events. In a recent investigation employing the HGF to model surprise about unexpected events, persons with autism learned more quickly about environmental volatility than controls without autism (Lawson *et al.*, 2017). However, at lower levels of the hierarchy, the tendency to believe that environmental instability is unstable resulted in smaller prediction errors (surprise) when events violated expectations. In other words, when the world is judged to be unstable and unpredictable, an agent differentiates less between expected and unexpected outcomes. This offers a similar but computationally distinct account of the stimulation-related learning changes described in a previous study (Seymour *et al.*, 2016), in which reduced positive and negative instrumental outcome sensitivity was reported as a consequence of neurostimulation. Similar to prior work, we found a positive relationship between model-based estimates of uncertainty and impulsivity (Averbeck *et al.*, 2013; Djamshidian *et al.*, 2012; FitzGerald *et al.*, 2015; Paliwal *et al.*, 2014). A plausible computational account of impulsivity is that high subjective uncertainty leads to lack of predictability and thus increases a tendency for short-term reward seeking and exploration.

We established that the ‘*standard*’ HGF best explained the gambling behaviour of our participants, in favour of a Rescorla-Wagner model or an ‘*uncertainty-driven*’ HGF. Importantly, the distinction between the standard and uncertainty-driven HGF models pertains only to the modelling of responses (the perceptual model is identical), in which the standard HGF employs a fixed decision temperature and the uncertainty-driven HGF a dynamic belief-to-response mapping based on online estimates of uncertainty (of beliefs about winning probability). These model comparison results suggest that our participants incorporate estimates of volatility into their prediction of reward probability but do not vary the stochasticity of their responses in response to these estimates. This is an interesting point of difference from the findings amongst younger, healthy males who completed a similar (albeit much longer) version of this task (Paliwal *et al.*, 2014) and future work will corroborate whether this finding of a static decision temperature is also observed amongst other neurodegenerative disorders.

In our participants, neurostimulation may interact with the physiology of the STN and alter the computations it implements. A tripartite functional organisation of the STN into limbic, associative and motor subregions is suggested by primate and human studies (Haynes and Haber, 2013; Lambert *et al.*, 2012), with electrode implantation targeted to the dorsolateral sensorimotor region to address motor symptoms of Parkinson’s disease (Wodarg *et al.*, 2012). Yet, the small size of the STN means that dispersion of electrical charge from a stimulating contact in this region could still modulate subthalamic regions with greater connectivity to fronto-striatal networks. The more ventral and medial the stimulating contact, the more likely these networks are to be affected by DBS. Previous investigations have suggested that the site of subthalamic stimulation can modulate cognitive (Hershey *et al.*, 2010) and psychiatric symptoms (Mallet *et al.*, 2007; Mosley *et al.*, 2018c; Welter *et al.*, 2014). How could STN-DBS modulate uncertainty? From a computational perspective, the STN has been considered to implement a ‘*delay*’ on cognitive-associative circuits in the basal ganglia, allowing more information to be gathered to guide the most appropriate behavioural policy, suppressing impulsive and potentially error-prone responding (Cavanagh *et al.*, 2011; Frank *et al.*, 2007). It is possible that by modulating the decision threshold, STN-DBS could alter the bound for evidence accumulation and thus uncertainty in the representation of the reward environment (Herz *et al.*, 2018; Pote *et al.*, 2016). Further work employing drift diffusion modelling to quantify rates of evidence accumulation and decision boundaries after STN-DBS may be illuminating, having previously helped to elucidate the mechanisms underlying hallucinations in Parkinson’s disease (O’Callaghan *et al.*, 2017). Further work is also required to determine if the site of stimulation affects the magnitude of changes in uncertainty estimation observed here and specifically if cognitive-associative or sensorimotor regions of the STN are most implicated in these shifts.

We did not observe a cross-sectional relationship between dopaminergic medication (expressed as LEDD) and uncertainty encoding, nor did we observe a longitudinal relationship between LEDD and self-reported impulsivity. However, there was a longitudinal relationship between changes in model parameter estimates and the maximum reduction in LEDD during longitudinal follow up. Specifically, the greater the increase in decision noise (the greater the decrease in *β*), the greater the postoperative reduction in LEDD. It is difficult to be certain about whether this is a causal relationship and it may be an epiphenomenon of effective subthalamic DBS: One of the benefits of the STN (as opposed to other surgical targets in DBS for Parkinson’s disease such as the internal segment of the globus pallidus) is that it allows for significant postoperative reduction in dopaminergic medication. Therefore, this apparent relationship could well be mediated by the effect of electrical stimulation, increasing indiscriminate responding and leading to a reduced requirement for dopaminergic therapy. Moreover, the finding that this relationship no longer held after the removal of an outlying participant decreases the confidence in this result.

There are likely to be fundamental differences in the computational operations subserved by the STN and dopamine in decision-making and impulsive behaviour. We have discussed the chronometric role of the STN is setting a decision bound and delaying impulsive choice, whereas dopamine is likely to have an essential role in reinforcement learning and reward evaluation (Abler *et al.*, 2006; Basar *et al.*, 2010; Daw *et al.*, 2006; Haber and Knutson, 2010; Kishida *et al.*, 2016; Schultz *et al.*, 1997; Tanaka *et al.*, 2008; Wittmann *et al.*, 2008). In a non-surgical population, persons with Parkinson’s disease withdrawn from medication display a characteristic impairment in reward learning and may show enhanced punishment sensitivity (Frank *et al.*, 2004). However, whilst dopamine replacement enhances the ability to learn from positive outcomes, learning from negative outcomes is impaired (Frank *et al.*, 2004). Thus, if postoperative LEDD reduction were a principal driver of a change in behaviour subsequent to DBS, then a selective impairment in positive outcome representation would be expected. However, from the HGF perspective, an agent with increased uncertainty at higher levels would be expected to show both decreased reward and punishment learning, as surprise to both positive and negative unexpected outcomes would be reduced (Lawson *et al.*, 2017). This suggests that LEDD changes may have a secondary role, but further careful experiments will be necessary to address this question. For example, the goal of this behavioural analysis was to relate a computational marker of uncertainty (over all trials) to impulsivity, but future neuroimaging investigations could model trial-wise positive and negative reward prediction errors and relate this to trial-wise brain activity. Participants could also be tested prior to STN-DBS ‘on’ and ‘off’ medication (although in our cohort, participants were too impaired by their movement disorder to tolerate this and a group of less severely-affected individuals would be required).

We did not observe significant correlations between behaviour or parameters inferred from slot machine play with other estimates of impulsivity including the excluded letter fluency task, the Hayling test and the delay discounting task. This reflects the multifaceted nature of impulsivity, which may implicate discrete subcortical and cortical regions and may evidence differential patterns of expression amongst impulsive endophenotypes (Nombela *et al.*, 2014; Robbins *et al.*, 2012). For example, the Hayling and ELF tasks are more commonly included amongst measures of task-switching and conflict interference, whilst the delay discounting task assesses impatience. Alternative paradigms may be required to capture participant-wise behaviour amongst these constructs.

We acknowledge methodological limitations of our investigation. The lack of a counterbalanced on-off stimulation design means that we cannot directly infer that stimulation underlies the observed changes in perceptual modelling observed in our participants (rather than, for example, practice effects or time). Specifically, for our participants, the pre-DBS session was the first time they had performed the task, and so changes in postoperative behaviour could also be attributable to greater familiarity with the task and perhaps the inherent volatility of the reward structure. However, we suggest that a strength of our longitudinal design is that it is more reflective of the natural clinical course taken by persons with Parkinson’s disease in the clinic. Moreover, our participants simply would not have tolerated an extended DBS washout and we hypothesise that the younger age of participants in the study of Seymour *et al* may have facilitated their crossover design (Seymour *et al.*, 2016). Nevertheless, it would be important to consider future experiments that could resolve this question, for example, selecting a cohort of younger Parkinson’s disease participants who could tolerate a washout of stimulation, or testing a cohort of participants without DBS twice, thirteen-weeks apart.

Unfortunately, in this study, we were unable to utilise estimates of meta-volatility in our analysis as *ϑ* could not be robustly recovered from simulated data. This failure to recover *ϑ* might result from the limited number of trials (100) completed by each participant, which limits the amount of information that can be gathered to update estimates of this higher-level HGF parameter from the population prior. Again, the disability of our participant cohort prohibited a greater number of trials, as employed in previous studies using this paradigm (Paliwal *et al.*, 2014), but this could be considered in future studies using younger or less severely-affected participants.

In summary, this study suggests that subjective estimates of uncertainty pertaining to environmental volatility and the stochasticity in belief-to-choice mapping change after subthalamic DBS for Parkinson’s disease and relate significantly to postoperative impulsivity. Increased estimates of environmental uncertainty (volatility) and increased noise in the decision process may contribute to impulsivity as a clinically relevant form of maladaptive behaviour. Uncertainty elevates the learning rate and suppresses top-down expectations, which may blunt error signalling in a series of trial-wise outcomes. Similarly, a consistent decision rule with regards to acting on an internal model of the world is important to make appropriate decisions based on what has been learned. We therefore posit a cognitive mechanism for the genesis of impulsive behaviour in this population. Finally, our results demonstrate that a naturalistic assessment of gambling behaviour in a virtual casino is useful for investigating impulsivity in Parkinson’s disease. The potential of our model to explain changes in impulsivity through game play could be most valuable in this disorder, given the significant, but poorly quantified risks relating to surgical (neurostimulation) and medical (dopamine agonist) treatments. If those at a higher risk of neuropsychiatric harm could be identified, this would improve the nature of treatment choice and informed consent and the effectiveness of clinical follow-up.

## 6 Supplementary Data

**Supplementary Table 8.1.** Additional neuropsychiatric assessment data pre- and post-DBS Significance: *** p<0.001, ** p<0.01, * p<0.05, where *p*-values are Holm-Bonferroni corrected for multiple comparisons with *α* = 0.05.

| Behavioural Measure | Pre-DBS | Post-DBS | Max Impairment | Pre- vs. Post-DBS |  |
| --- | --- | --- | --- | --- | --- |
|  | *Mean (SD), Median (Range)* |  |  | *t-stat* | *p-value* |
| **BIS Attentional** | 16.1 ($\pm 3.2$),<br>16 (10 - 23) | 15.0 ($\pm 3.4$),<br>15 (8 - 22) | 0.39 ($\pm 2.8$),<br>0 (-5 - 5) | 2.64 | 0.06 |
| **BIS Non-Planning** | 23.0 ($\pm 4.1$),<br>23 (14 - 33) | 22.4 ($\pm 5.2$),<br>22 (15 - 36) | 2.3 ($\pm 3.7$),<br>2 (-6 - 9) | 1.09 | 0.84 |
| **BIS Motor** | 21.2 ($\pm 3.3$),<br>21 (14 - 29) | 20.3 ($\pm 3.4$),<br>21 (13 - 30) | 1.2 ($\pm 3.5$),<br>1 (-7 - 14) | 1.87 | 0.28 |
| **Apathy Scale** | 11.5 ($\pm 5.7$),<br>11 (1 - 26) | 11.3 ($\pm 5.5$),<br>11 (1 - 23) | 2.8 ($\pm 4.2$),<br>2 (-3 - 14) | 0.30 | 1.54 |
| **Empathy Quotient** | 40.7 ($\pm 13.1$),<br>39.5 (16 - 68) | 40.4 ($\pm 14.3$),<br>36 (14 - 68) | -4.7 ($\pm 7.9$),<br>-5 (-26 - 13) | 0.27 | 1.54 |
| **Geriatric Anxiety Inventory** | 4.9 ($\pm 4.3$),<br>5 (0 - 17) | 3.0 ($\pm 4.0$),<br>2 (0 - 20) | 0.55 ($\pm 3.4$),<br>0 (-5 - 11) | 3.62 | <0.001* |

**Supplementary Figure 8.1.**
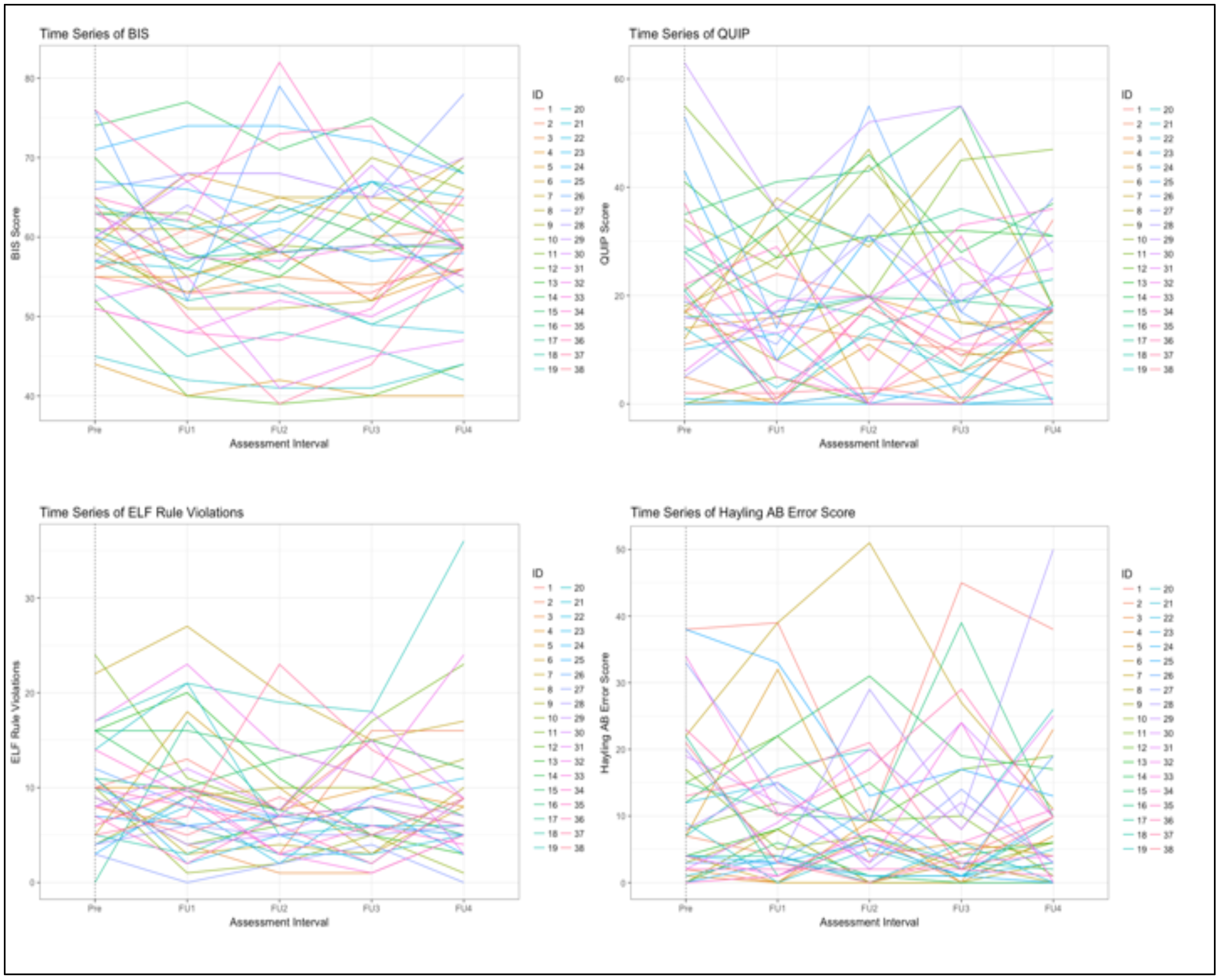
Heterogeneity in participant-wise trajectories. The participant-wise trajectories of key variables were inspected across all intervals in the investigation. Representative figures are presented below. Although mean changes were not significantly different at postoperative assessments, there was considerable inter-individual heterogeneity in postoperative course. BIS = Barratt Impulsiveness Scale, QUIP = Questionnaire for Impulsive-Compulsive Disorders in PD, ELF = Excluded Letter Fluency. Pre = Pre-DBS, FU1 = 2-weeks post-DBS, FU2 = 6-weeks post-DBS, FU3 = 13-weeks post-DBS, FU4 = 26-weeks post-DBS.

**Supplementary Table 8.2.** Pre-DBS slot machine behaviour and pre-DBS BIS subscales *b* values are standardized regression coefficients. Significance: *** p<0.001, ** p<0.01, * p<0.05, where *p*-values are Holm-Bonferroni corrected for multiple comparisons with *α* = 0.05. ^ Indicates significant t-statistics, Holm-Bonferroni corrected for multiple comparisons.

|  | Independent Variables (*b*) |  |  |  |  |  |  |
| --- | --- | --- | --- | --- | --- | --- | --- |
| Dependent Variables | Bet Size | Machine Switch | Double-up | BDI | $R^2$ | F-stat | *p*-value |
| BIS Non-Planning | 0.012 | -0.028 | -0.056 | 0.343 | 0.179 | 1.801 | 0.152 |
| BIS Motor | -0.003 | 0.266 | -0.046 | 0.138 | 0.094 | 0.858 | 0.499 |
| BIS Attentional | 0.025^ | 0.100 | 0.025 | 3.314 | 0.332 | 4.094 | 0.008** |

**Supplementary Table 8.3.** Post-DBS slot machine behaviour and post-DBS BIS subscales *b* values are standardized regression coefficients. Significance: *** p<0.001, ** p<0.01, * p<0.05, where *p*-values are Holm-Bonferroni corrected for multiple comparisons with *α* = 0.05. ^ Indicates significant t-statistics, Holm-Bonferroni corrected for multiple comparisons.

|  | Independent Variables (*b*) |  |  |  |  |  |  |
| --- | --- | --- | --- | --- | --- | --- | --- |
| Dependent Variables | Bet Size | Machine Switch | Double-up | BDI | $R^2$ | F-stat | *p*-value |
| BIS Non-Planning | 0.006 | -0.237 | 0.031 | 0.439 | 0.303 | 3.588 | 0.016* |
| BIS Motor | -0.004 | 0.467 | 0.115 | 0.026 | 0.250 | 2.753 | 0.044* |
| BIS Attentional | 0.016^ | -0.101 | 0.123^ | 0.254 | 0.496 | 8.123 | <0.001*** |

**Supplementary Table 8.4.**
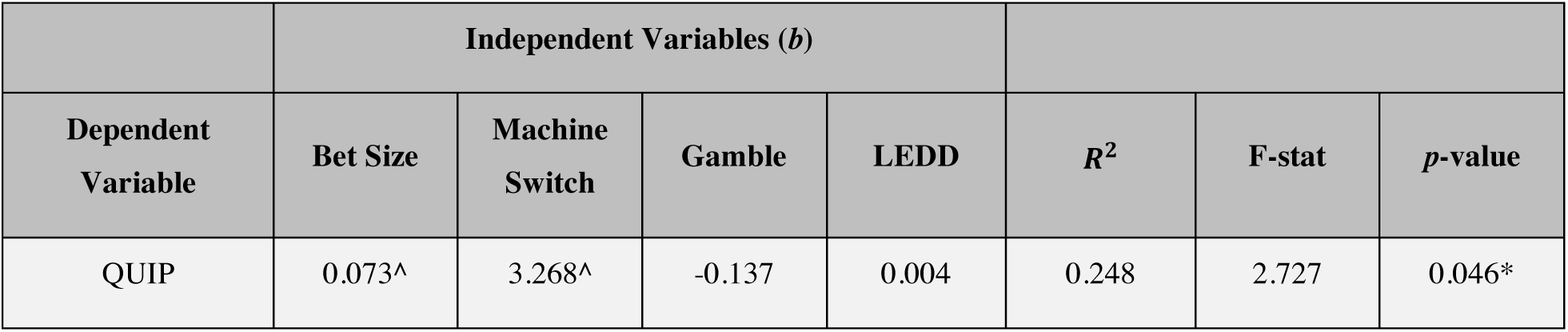
Post-DBS gambling behaviour and post-DBS QUIP-RS The QUIP and LEDD correlated strongly at both time points (*ρ*_*pre*_=0.42, *p*=0.008; *ρ*_*post*_=0.44, *p*=0.005), with LEDD decreasing significantly post-DBS. LEDD was therefore included as a covariate when regressing against QUIP-RS total scores, in order to explain the remaining variance in QUIP due to gambling behaviour. This is consistent with a previous association of dopaminergic medication with compulsive behavioural disorders in PD (Weintraub *et al.*, 2010). Higher bets (*t*_(37)_=2.057, *p*=0.048) and more frequent machine switches (*t*_(37)_=3.268, *p*=0.016) corresponded with higher QUIP scores. *b* values are standardized regression coefficients. Significance: *** p<0.001, ** p<0.01, * p<0.05, where *p*-values are Holm-Bonferroni corrected for multiple comparisons with *α* = 0.05. ^ Indicates significant t-statistics, Holm-Bonferroni corrected for multiple comparisons.

**Supplementary Table 8.5.** Pre-DBS model parameters and pre-DBS BIS subscales *b* values are standardized regression coefficients. Significance: *** p<0.001, ** p<0.01, * p<0.05, where *p*-values are Holm-Bonferroni corrected for multiple comparisons with *α* = 0.05. ^ Indicates significant t-statistics, Holm-Bonferroni corrected for multiple comparisons.

| | Independent Variables ($b$) | | | | | |
| --- | --- | --- | --- | --- | --- | --- |
| Dependent Variables | $\omega$ | $\beta$ | BDI | $R^2$ | F-stat | $p$-value |
| BIS Non-Planning | -3.268 | -0.056 | 0.326 | 0.172 | 2.351 | 0.090 |
| BIS Motor | -1.222 | -0.088 | 0.085 | 0.020 | 0.226 | 0.877 |
| BIS Attentional | -0.817 | -0.225 | 0.318 | 0.226 | 3.314 | 0.031* |

**Supplementary Table 8.6.**
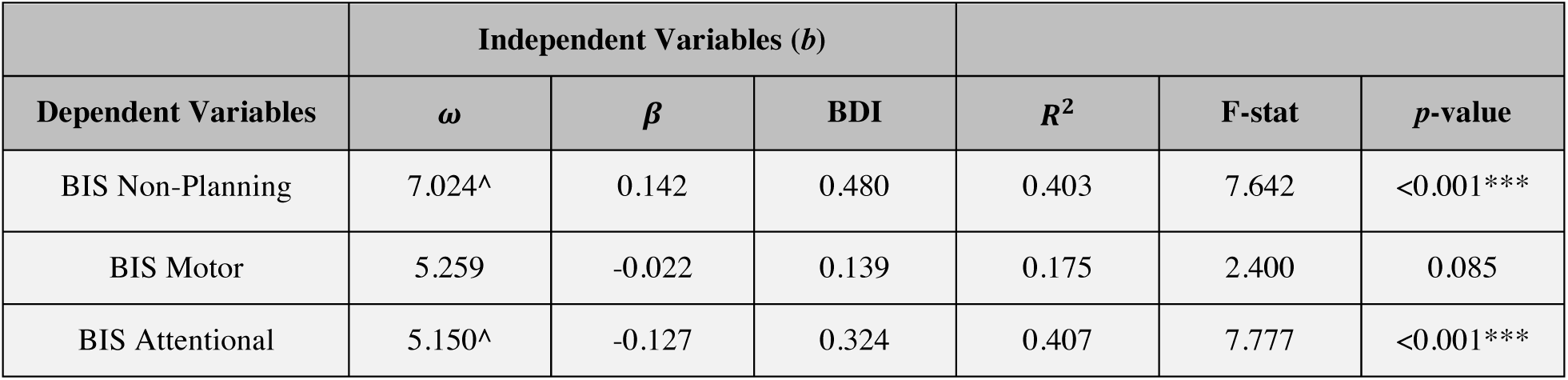
Post-DBS model parameters and post-DBS BIS subscales *b* values are standardized regression coefficients. Significance: *** p<0.001, ** p<0.01, * p<0.05, where *p*-values are Holm-Bonferroni corrected for multiple comparisons with *α* = 0.05. ^ Indicates significant t-statistics, Holm-Bonferroni corrected for multiple comparisons.

| | Independent Variables ($b$) | | | | | |
| --- | --- | --- | --- | --- | --- | --- |
| Dependent Variables | $\omega$ | $\beta$ | BDI | $R^2$ | F-stat | $p$-value |
| BIS Non-Planning | 7.024^ | 0.142 | 0.480 | 0.403 | 7.642 | <0.001*** |
| BIS Motor | 5.259 | -0.022 | 0.139 | 0.175 | 2.400 | 0.085 |
| BIS Attentional | 5.150^ | -0.127 | 0.324 | 0.407 | 7.777 | <0.001*** |

**Supplementary Table 8.7.**
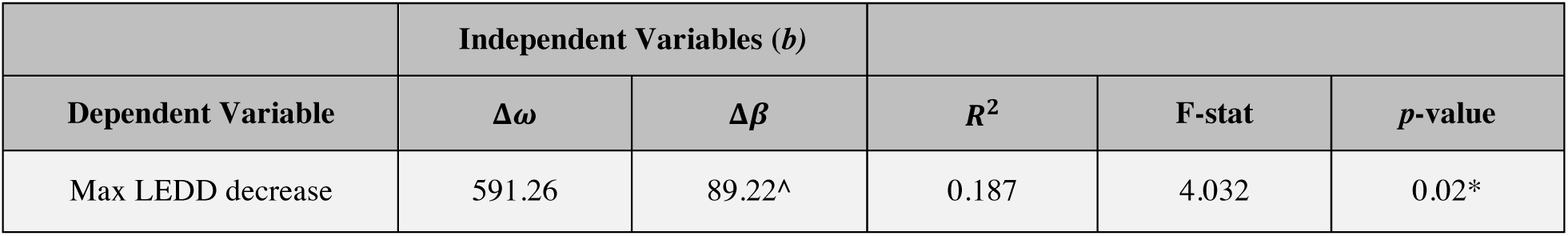
Max LEDD reduction and change in model parameters *b* values are standardized regression coefficients. Significance: *** p<0.001, ** p<0.01, * p<0.05, where *p*-values are Holm-Bonferroni corrected for multiple comparisons with *α* = 0.05. ^ Indicates significant t-statistics, Holm-Bonferroni corrected for multiple comparisons. *Changes in ω and β are calculated as the pre-DBS parameter estimate value minus the post-DBS parameter estimate value. Max LEDD decrease refers to the maximum postoperative reduction of dopaminergic medication across all measurement intervals subsequent to DBS.*

**Supplementary Figure 2.**
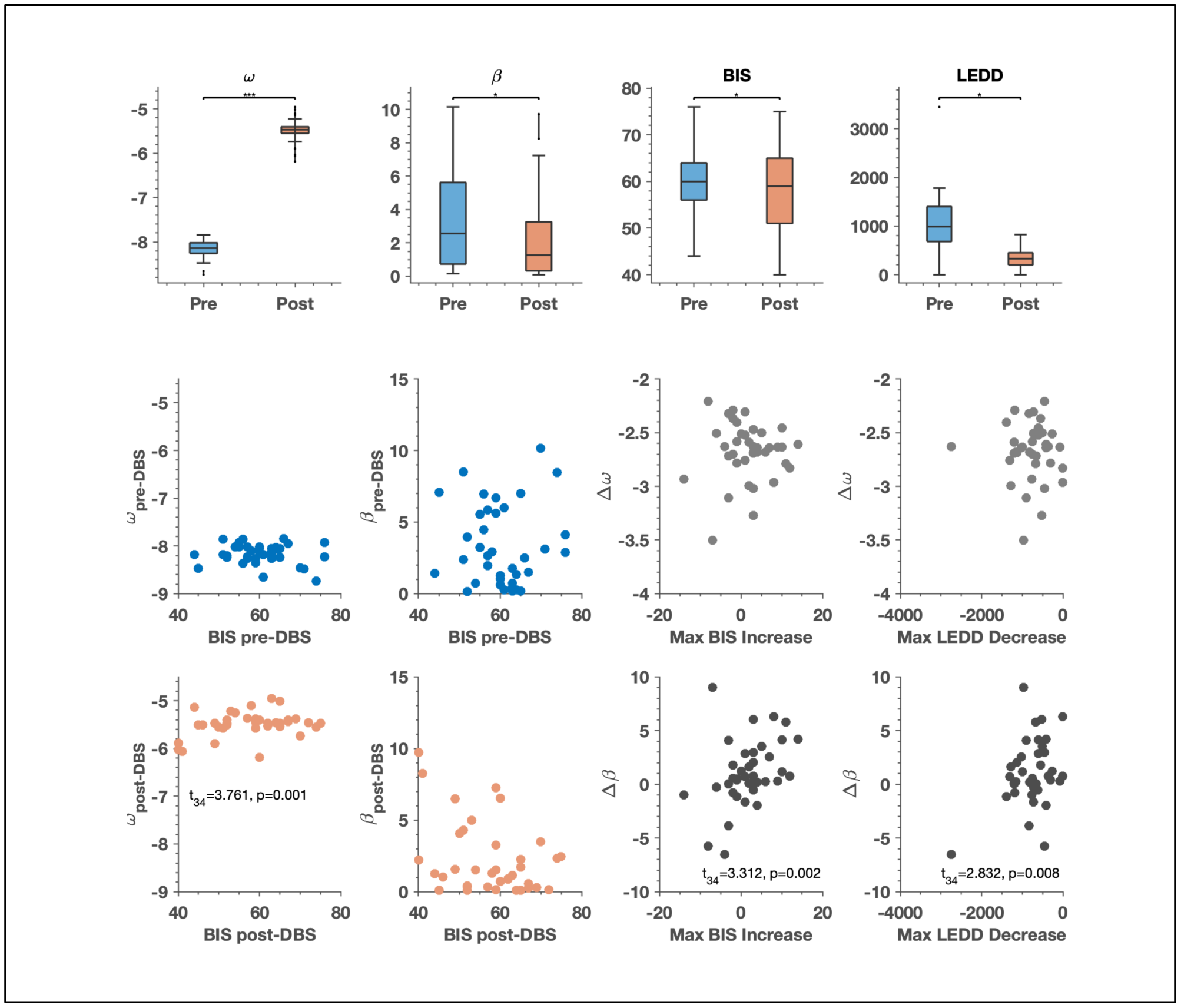
Computational model parameters, pre-and post-DBS. Shown in the figure above is a version of Figure 8.7 in the main manuscript, with one change. Here, we have included the outlier in assessing the relationship between the pre-to-post operative change in *β* with the Max LEDD Decrease. As can be seen here, the outlier is driving the statistical relationship between these variables. The full figure is replicated here to provide the context of these analyses.

## 7 Acknowledgements

The authors gratefully acknowledge the commitment of patients and caregivers who contributed their time to this study. The authors acknowledge the ongoing support of St Andrew’s War Memorial Hospital and the Herston Imaging Research Facility.

## 8 Data availability statement

The HGF toolbox is part of the open source TAPAS software and available for download at http://www.translationalneuromodeling.org/tapas. The gambling paradigm is provided for download on a git repository at https://github.com/saeepaliwal/breakspear_slot_machine.git. The analysis pipeline is provided at https://github.com/saeepaliwal/dbs_pd_analysis_pipeline.git. A de-identified data set containing neuropsychiatric assessment and gambling data can be provided by Dr Philip Mosley on application, subject to institutional review board approval.

## 10 Funding agencies

PEM was supported by an early career fellowship from the Queensland government’s ‘Advance Queensland’ initiative, a Royal Brisbane & Woman’s Hospital Foundation Research Grant and a young investigator grant from the Royal Australian and New Zealand College of Psychiatrists. He received an unrestricted educational grant from Medtronic. MB was supported by the National Health and Medical Research Council (118153, 10371296, 1095227) and the Australian Research Council (CE140100007). KES was supported by the University of Zurich and the René and Susanne Braginsky Foundation.

## 11 Author Contribution Statement

Paliwal: Task design, statistical analysis, writing of first draft of manuscript*

Mosley: Data collection, statistical analysis, writing the first draft of the manuscript *

Breakspear: Critical comments on study design and manuscript

Coyne: Supervision of data collection, critical comments on manuscript

Silburn: Supervision of data collection, critical comments on manuscript

Aponte: Collaborated on model inversion and cross validation

Mathys: Collaborated on model inversion and cross validation

Stephan: Task design, supervision of data analysis, contributions to manuscript writing

* These authors contributed equally to the work

